# Life on leaves left a lasting mark on the *Methylobacterium* core genome

**DOI:** 10.64898/2026.11.02.763778

**Authors:** Jean-Baptiste Leducq, David Sneddon, Alexander B. Alleman, Pratibha Sanjenbam, Jack M. Sullivan, Deepa Agashe, Christopher J. Marx

**Affiliations:** Université Laval, Québec City, QC, Canada; University of Idaho, Moscow, ID, USA; Oregon State University, Corvallis, OR, USA; Western Colorado University, Gunnison, CO, USA; National Centre for Biological Sciences, Bengaluru, India

**Keywords:** Phylogenetic discordance, Core genome evolution, *Methylobacterium*, Phyllosphere adaptation, Anoxygenic photosynthesis, Carotenoid biosynthesis, Horizontal gene transfer, Phylogenomics

## Abstract

The extent to which core genomes reflect strictly vertical inheritance remains debated, particularly in ecologically diverse bacterial lineages. Here, we investigate the evolution of the core genome of *Methylobacterium*, a major leaf-associated bacterial genus, to assess how evolutionary signals vary across genes and whether they relate to functions involved in adaptation to life at the surface of plant leaves (the phyllosphere). Identifying 1,099 protein-coding core genes across 293 genomes, we combined gene tree comparisons, phylogenetic network reconstruction, nucleotide polymorphism patterns, and functional enrichment analyses. We found that most core genes show some deviation from the consensus phylogenetic tree. However, this discordance is not randomly distributed. Across all analytical approaches, genes showing the strongest deviations are consistently associated with functions linked to life in the phyllosphere, including anoxygenic photosynthesis, protection against UV and osmotic stress. These genes are often grouped into conserved clusters in the genome. In particular, a large cluster involved in anoxygenic photosynthesis, including bacteriochlorophyll and carotenoid biosynthesis, shows strong variation in its evolutionary history and organization across lineages, despite maintaining a conserved core structure. Similar patterns of phylogenetic incongruence and gene organization are observed across related Hyphomicrobiales inhabiting the phyllosphere, suggesting that these functions extend beyond *Methylobacterium*. Together, these results indicate that evolutionary heterogeneity within the core genome is concentrated in genes involved in key ecological functions. While alternative explanations cannot be excluded, these patterns are consistent with horizontal gene transfer contributing to the diversification of traits important for survival and adaptation in phyllosphere bacterial communities.

## Introduction

The phyllosphere, defined as the aerial surfaces of plants, represents one of the largest microbial habitats on Earth and hosts diverse communities of prokaryotes and fungi that play key ecological roles in plant-associated ecosystems (Lindow and Brandl 2003; Vorholt 2012). Microorganisms colonizing leaf surfaces contribute to plant growth, stress tolerance, and ecosystem functioning, but must also cope with strong and rapidly fluctuating environmental conditions characteristic of this habitat (Vorholt 2012; Remus-Emsermann and Schlechter 2018). Among these constraints, exposure to solar radiation, in particular ultraviolet (UV) radiation, desiccation, and nutrient limitation, constitutes major selective pressures shaping microbial community structure and function on leaves (Jacobs and Sundin 2001; Gamit et al. 2025). Survival in the phyllosphere therefore relies on a suite of physiological and metabolic traits that enable microorganisms to mitigate oxidative stress, maintain cellular integrity, and exploit transient resources. Many of these traits have been characterized at the functional level, including mechanisms related to stress tolerance and resource acquisition, yet their distribution across bacterial lineages and the evolutionary processes underlying their diversification remain incompletely understood (Delmotte et al. 2009; Vorholt 2012).

In bacteria, genome evolution is shaped by the interplay between vertical inheritance and horizontal genetic exchange, resulting in mosaic genomes in which different genes have distinct evolutionary histories. This property complicates the reconstruction of a single, consistent phylogenetic history and can generate substantial discordance between gene trees and species relationships (Maddison 1997; Degnan and Rosenberg 2009). While such phylogenetic discordance may arise from multiple biological and methodological processes, including incomplete lineage sorting (ILS) and stochastic variation in phylogenetic inference, it is also commonly interpreted as a hallmark of horizontal gene transfer (HGT) in microbial systems. Distinguishing between these alternative explanations remains challenging, particularly at the scale of large comparative genomic datasets. Core genes, defined as the set of genes shared by individuals within a given bacterial group, are often assumed to follow a predominantly vertical evolutionary history, as they typically encode essential cellular functions under strong purifying selection. For this reason, they are commonly used to infer consensus phylogenies (Chung et al. 2018). However, growing evidence indicates that horizontal gene transfer can also affect core genes, albeit less frequently than accessory genes (Koonin et al. 2001; Polz et al. 2013). In addition, the composition and size of the core genome depends strongly on the phylogenetic scale considered: closely related lineages tend to share a larger set of genes than more distantly related taxa (Park et al. 2019). As a result, increasing phylogenetic resolution, and thus the number of core genes, can improve the statistical support for phylogenetic trees, on one hand, but can also increase the likelihood of capturing genes exhibiting phylogenetic discordance, on the other.

*Methylobacterium* (order Hyphomicrobiales) is a genus of widespread plant-associated methylotrophic bacteria that are frequently found on leaf surfaces and can influence plant physiology through promoting host growth, mitigating stress, modulating photosynthesis, and protecting against UV damage (Knief et al. 2010; Sanjenbam et al. 2022). *Methylobacterium* species can utilize methanol released by plants as a carbon and energy source (Delmotte et al. 2009). Members of the genus also produce carotenoids that contribute to protection against oxidative and UV-induced stress (Van Dien et al. 2003; Mo et al. 2023) and are capable of aerobic anoxygenic photosynthesis through bacteriochlorophyll-based phototrophic systems (Stiefel et al. 2013; Ma et al. 2024): key adaptations for survival on leaf surfaces. Recent phylogenomic analyses have revealed that *Methylobacterium* is structured into four deeply divergent phylogenetic groups (thereafter phylogroups A, B, C, D) that differ in genome content, architecture, and ecological distribution (Leducq, Sneddon, et al. 2022). Notably, the most ancestral phylogroup of *Methylobacterium* (C) is primarily soil-or root-associated, whereas the later-diverging phylogroups (A, B, D) are predominantly found in the phyllosphere, suggesting a secondary adaptation to aerial plant surfaces (Lee et al. 2022; Alleman et al. 2025). More broadly, members of the Hyphomicrobiales represent a diverse and abundant component of phyllosphere microbial communities, although their full diversity has only recently begun to be characterized through phylogenomic and metagenomic approaches (Leducq et al. 2026).

Several *Methylobacterium* lineages are frequently observed co-occurring within the same plant hosts and environments, suggesting that closely related but genetically distinct populations may share ecological niches at fine spatial scales (Knief et al. 2010; Leducq, Seyer-Lamontagne, et al. 2022; Sanjenbam and Agashe 2024). Such ecological overlap may create opportunities for genetic exchange among lineages during diversification, potentially contributing to the maintenance or reshaping of adaptive traits (Polz et al. 2013; Woods et al. 2020; Coyte et al. 2022). At the same time, phylogenetic relationships among phyllosphere-associated *Methylobacterium* phylogroups A, B, and D remain partially unresolved, consistent with either rapid lineage divergence or more complex evolutionary processes involving gene exchange (Leducq, Sneddon, et al. 2022).

In this study, we investigated the evolutionary dynamics of the *Methylobacterium* core genome to identify signatures of phylogenetic discordance and assess whether specific subsets of genes exhibit distinct evolutionary patterns. We first generated a dataset of 1,099 protein-encoding core genes across 293 *Methylobacterium* genomes, then combined multiple complementary approaches, including gene tree–consensus tree comparisons, evolutionary network analyses, nucleotide polymorphism patterns, and functional enrichment tests. We provide strong evidence that a subset of core genes displays unusually high levels of phylogenetic discordance compared to the genomic background. While our inferences are based on patterns of discordance and therefore remain compatible with several underlying processes, the systematic detection of those patterns for genes involved in anoxygenic photosynthesis and carotenoid biosynthesis through independent methods is consistent with a scenario in which horizontal genetic exchanges have contributed to the diversification of those key functions in *Methylobacterium*, and potentially in related Hyphomicrobiales genera inhabiting the phyllosphere. More generally, our results suggest that the evolutionary trajectories of genes involved in phyllosphere adaptation may be more dynamic than those of the core genome as a whole.

## Results

### Methylobacterium core genome definition

We compiled a dataset of 293 *Methylobacterium* genomes for core genome definition, including 189 previously annotated genomes and 104 newly incorporated assemblies (Table S1). The latter comprised 79 new rice-associated genomes from this study (Table S2; Figure S1), 7 soybean-associated genomes, and 18 recently published genomes, including representatives of newly described species (see Method).

A total of 9,259 unique gene annotations from RAST were identified across the 293 genomes. Applying relaxed thresholds for core gene definition within phylogroups initially yielded 1,419 candidate core genes based on the 189 previously annotated genomes. After inclusion of the 104 new genomes and correction for artefactual gene duplications caused by assembly fragmentation, 1,122 core genes were retained, including 1,099 protein-encoding genes (PEGs) and 23 RNA genes (Table S3).

The number of core genes retrieved per genome varied substantially (mean = 1,093; range = 928–1,118; Table S1). This variation was primarily associated with genome assembly quality, with lower N50 assemblies showing reduced recovery of core genes, while no clear effect of phylogroup identity was observed (Figure S2). Only the 1,099 PEGs were retained for downstream phylogenomic analyses.

### Methylobacterium species and lineage delimitation, and phylogenomic analysis

We classified *Methylobacterium* species and lineages based on ANI (whole genome Average Nucleotide Identity) and phylogenomic reconstruction (core genome). Pairwise ANI values across the 293 genomes showed a multimodal distribution (Figure S3) and were used to delineate species using a ≥95% threshold. This resulted in the identification of 116 *Methylobacterium* species (1-29 genomes per species; Figure 1A; Table S4). Species assignments of the 189 previously analyzed *Methylobacterium* assemblies were mostly consistent with the 104 species from our previous study (Leducq, Sneddon, et al. 2022), with the exception of 4 species fusions (*M. sp. 015* with *M. bullatum*, *M. sp. 033* with *M. litchi*, *M. sp. 042* with *M. sp. 043*; *M. sp. 045* with *M. alsaeris*). Of the 79 new *Methylobacterium* isolate assemblies from rice (Sanjenbam et al. 2020), 65 were assigned to previously described or candidate species (*M. fujisawaense*: n=4; *M. radiotolerans*: n=22; *M. sp. 028*: n=8; *M. aerolatum*: n=1; *M. komagatae*: n=3; *M. thiocyanatum*: n= 1; *M. zatmanii*: n=2; *M. salsuginis*: n=13; *M. suomiense*: n=3; *M. ajmalii*: n=4; *M. indicum*: n=3; *M. gossipiicola*: n=1) and 14 were assigned to new candidate species (*M. sp. 046-049* and *051-053*). The 7 soybean leaf-associated genome assemblies (Govindaraju et al. 2025) were assigned to *M. extorquens* (n=4) and *M. sp 036* (n=3). Other assemblies (n=10) were assigned to *M. sp. 021* (n=2), *M. sp. 027* (n=1), *M. extorquens* (n=4), *M. organophilum* (n=1), *M. sp. 009* (n=1) and to a new candidate species (*M. sp. 050*; n=1). No genome from our dataset was assigned to recently described species *M. flocculans*, *M. oryzisoli*, *M. carpenticola*, *M. guangdongense*, *M. nigriterrae*, and *M. subtropicum* (Table S1).

**Figure 1:**
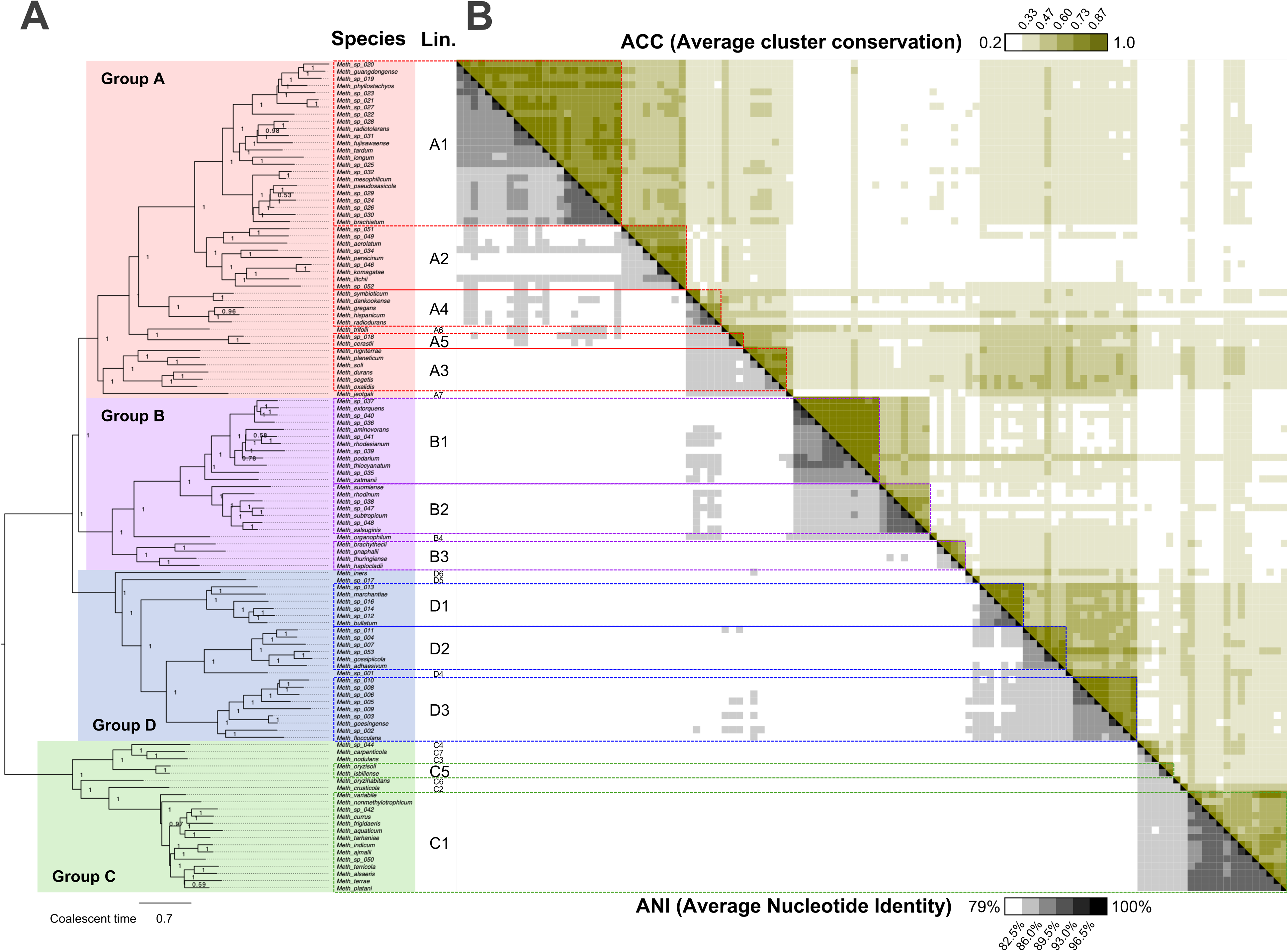
*Methylobacterium* species, lineage, and phylogroup definition is supported by core-genome phylogeny, genome-wide average nucleotide identity (ANI), and average gene cluster conservation (ACC). A) *Methylobacterium* species tree inferred with ASTER (116 species defined from 293 genomes grouped if ANI ≥95%; 1-29 genomes per species), combining 1,099 protein-encoding core gene phylogenies (ML trees individually computed with RAxML, 1,000 replicates). The four phylogroups (filled frames: A: red; B: purple; D: blue; C: green) and the 24 embedded lineages (open frames) are strongly supported by monophyly (local posterior probability = 1). The tree is rooted on phylogroup C. B) Heatmap showing pairwise ANI (below diagonal; grey scale) and ACC (above diagonal; yellow scale) among species (averaged among genomes from the same species). ANI was computed with FastANI. Cluster conservation indicates the proportion of same pairs of core genes located within 25 kb in two different genomes.

Phylogenomic analyses based on the 1,099 core PEGs produced both a consensus lineage tree (each genome treated as an independent evolutionary unit) and a species tree (genomes grouped by ANI-defined species), inferred under the multispecies coalescent model. The species tree resolved 116 species organized within 24 lineages and four phylogroups (A, B, C, D; Figure 1A; Figure S4). All species were recovered as monophyletic with maximal node support (local posterior probability = 1). Lineages were defined as monophyletic clades in the consensus lineage tree with maximal node support (local posterior probability = 1; Figure 1A, Figure S4) and corresponded to intermediate ANI values below the species threshold (ANI < 95%) but above phylogroup divergence (ANI ≥ 84% (Alleman et al. 2025)). Across lineages, ANI values used for delineation ranged approximately between 88% and 95%, with no single universal threshold. Lower bounds of ANI delimitation varied among lineages, typically between 84% and 88% (Figure 1B; Figure S3; Table S4).

### Gene cluster definition based on coevolution (pRF) and shared KEGG pathways (pKEGG)

To determine whether physically linked genes also share similar evolutionary histories in *Methylobacterium*, we first evaluated the relationship between genomic distance along the complete *M. oryzae* CBMB20 reference genome (phyllosphere-associated strain with the best genome assembly), phylogenetic similarity, and functional similarity among pairs of core PEGs based on KEGG pathways. KEGG pathway annotation identified 134 KEGG pathways across 649 of the 1,099 core PEGs, while no KEGG pathway was retrieved for the remaining 450 genes (Tables S3 and S5). Across the 1,099 protein-encoding core genes, we analyzed all 113,169 possible pairwise comparisons. Pairwise phylogenetic dissimilarity (*pRF*) and the probability that two genes shared at least one KEGG pathway (*pKEGG*) both significantly varied as a function of pairwise genomic distance among PEGs (*pDist*), with a breakpoint detected in slopes of both non-linear regressions at 8.5 and 31.5 kb, respectively (Davies test: p<<0.001; Figure S5A-B). Slopes of segmented linear regressions were stronger below than above breakpoints for both *pRF* (β *=* 4.44 [95% CI: 3.52 ; 5.37] and 0.22 [0.17 ; 0.27], respectively) and *pKEGG* (β *=* -0.103 [-0.127 ; -0.080] and 0.011 [0.008 ; 0.014], respectively), indicating that genes located within a few kilobases tend to share stronger functional and evolutionary associations than more distant genes. Because the relationships weakened with distance and because genome assemblies remained fragmented for many isolates, we defined gene clusters as genes located within 25 kb of one another for all downstream analyses.

To quantify whether the organization of co-localized genes (thereafter clusters) remained evolutionarily conserved despite extensive genomic rearrangements across the genus, we quantified average cluster conservation (ACC) across genomes and species. ACC closely followed the phylogenomic structure of the genus (Figure 1B). As expected, genomes belonging to the same species (ACC mean: 0.99 [range: 0.90 – 1.00]) or lineage (0.83 [0.68 – 0.94]) consistently shared higher proportions of conserved core-gene colocalization than genomes from distinct lineages (0.49 [0.31-0.83]) or phylogroups (0.37 [0.23-0.64]; Table S4; Figure S6). Overall, average ACC and ANI values measured within and among *Methylobacterium* species were significantly correlated (R^2^ = 0.71; p<0.001; Figure S6).

We next tested whether conservation of gene colocalization across lineages (CCL) was associated with phylogenetic similarity among genes. Across all pairwise comparisons, genes that colocalized in a larger number of lineages displayed lower pairwise phylogenetic dissimilarity (*pRF*), indicating more similar evolutionary histories (Figure S5C). This negative relationship was significant both when considering the complete set of 113,169 pairwise comparisons (β = -5.90 ± 0.35; t-test: p <0.001) and when averaging data across 50 CCL bins to reduce biases caused by the predominance of unrelated gene pairs (β = -6.06 ± 1.01; t-test: p <0.001; Figure S5C). These results suggested that, even across lineages, co-localized genes often moved together, resulting in shared evolutionary histories.

### Phylogenetic network approach and phylogenetic incongruence enrichment in KEGG pathways

To characterize phylogenetic incongruence among *Methylobacterium* lineages, we reconstructed an evolutionary network from the phylogenetic trees of a subset of 970 core PEGs (tree with at least two missing lineages removed) simplified at the lineage level, hence focusing on gene swaps prior lineage radiation (Figure 2A; Table S6). Considering nodes of the ASTER species tree, simplified at the lineage level, as nodes for our network reconstruction, we predicted 891 possible edges in the network, corresponding to non-redundant pairwise combinations among those nodes (thereafter partitions). Of these predicted partitions, 414 occurred in at least one of the 970 core PEG trees, including the 24 lineages (lineage partitions), 21 partitions corresponding to the consensus tree (consensus partitions), and 369 alternative partitions (out of the 846 predicted; 43.6%), i.e. partition that could be interpretated as a single branch swap compared to the consensus tree, hence putative gene exchanges during *Methylobacterium* evolution. In addition, we observed 1,424 alternative partitions that were not predicted, because they could not be interpreted as a single branch swap and were excluded from further analyses for simplification. The 21 consensus partitions were dominant, occurring in average in 466.6 ± 231.7 trees. The root of the consensus tree supporting phylogroup ancestral configurations AB/CD was observed in 77 trees but challenged by alternative phylogroup partitions AD/BC (177 trees) and BD/AC (76 trees).

**Figure 2.**
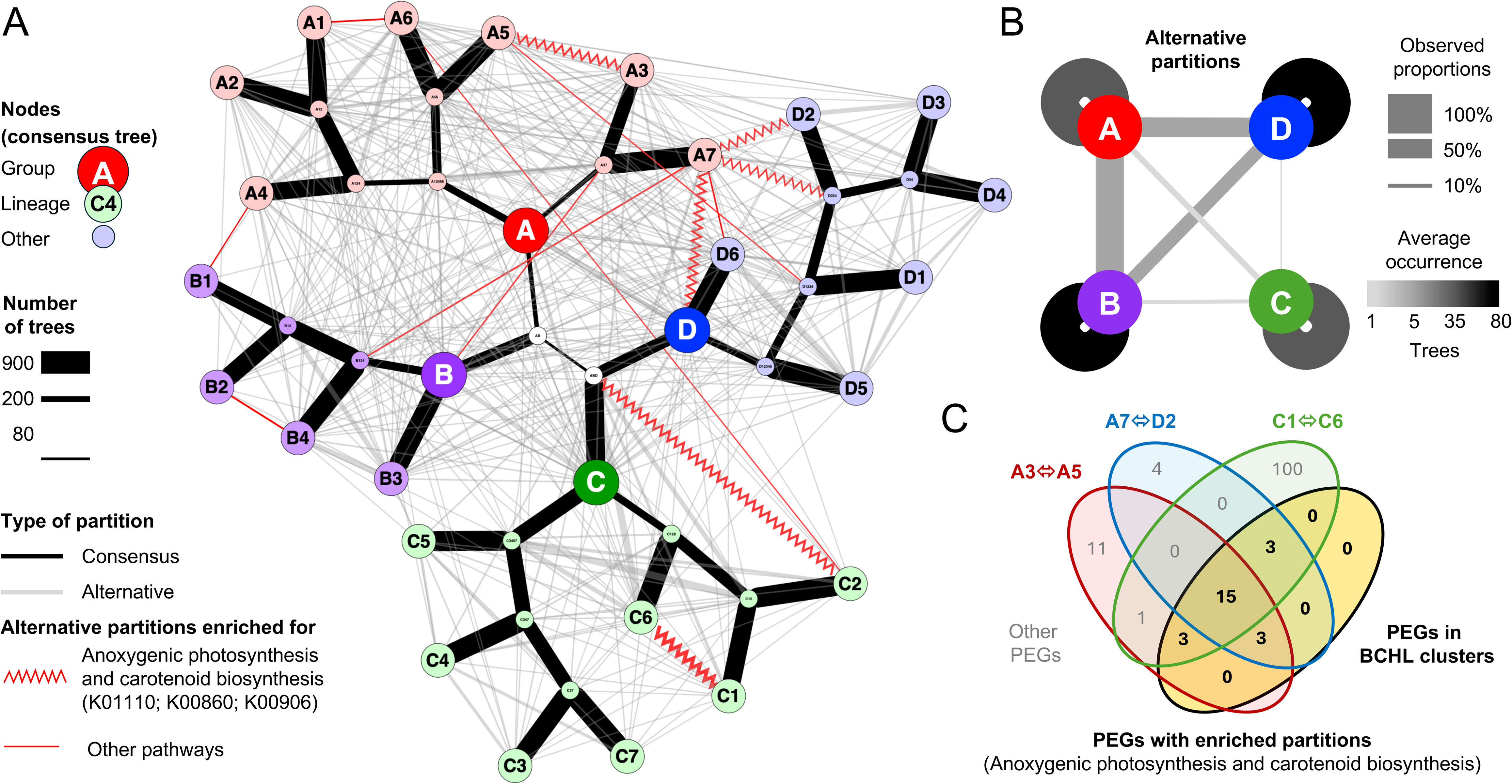
Evolutionary network of the *Methylobacterium* core genome suggests extensive gene exchanges among lineages. A) The network was reconstructed wby combining nodes and nested branches (partitions) of phylogenetic trees of 970 core PEGs. Edges are colored according to partition congruence with the consensus tree (black), incongruence (grey), and significantly enriched for KEGG pathways (01110: Biosynthesis of secondary metabolites; 00906: porphyrin metabolism; 00860: carotenoid biosynthesis; red; p < 0.01; 10,000 permutations; BH correction). Edge width is proportional to the number of trees supporting each partition. B) Gene swaps are more frequent within phylogroups and among phylogroups A, B and D than with phylogroup C. Line color intensity is proportional to the mean number of PEG trees supporting alternative partitions in each category, while line width is proportional to the proportion of predicted alternative partitions that were observed. C) Venn diagram showing the number of PEGs overlapping among the A3–A5 (red), A7–D2 (blue), and C1–C6 (green) alternative partitions enriched for KEGG pathways K01110, K00860, and K00906 (anoxygenic photosynthesis and carotenoid biosynthesis) and the BCHL cluster (yellow; bold).

We determined whether alternative partitions were enriched for branch swaps within phylogroups compared to swaps among phylogroups (Figure 2B, Table S6). Focusing on 698 predicted alternative partitions corresponding to swaps occurring after phylogroup radiation, 278 were observed in at least one PEG tree (39.8%). This proportion was significantly higher for partitions supporting swaps within phylogroups (107 observed out of 110 predicted) than for swaps between phylogroups A, B or D (138 out of 258; proportion test: p < 0.001). Both were significantly higher than the proportion of observed partitions supporting swaps between phylogroup C and either phylogroups A, B or D (33 out of 336; proportion test: p < 0.001). In average, partitions supporting branch swaps between phylogroup C and either phylogroups A, B or D (1.2 ± 0.4 trees) or among phylogroups A, B or D (4.1 ± 4.8 trees) were significantly rarer than partitions supporting branch swaps within phylogroups (45.0 ± 60.2 trees; Tukey test: p< 0.001).

Permutation analyses of KEGGs among PEGs showed that 14 partitions were significantly enriched for 13 KEGG pathways (100,000 permutations; p<0.05; Figure 2A; Table S6). All enriched partitions corresponded to alternative partitions, supporting gene swaps between lineages either within phylogroups (n=4), among phylogroups (n=7), or undetermined (n=3). Six partitions were associated with three overlapping KEGG pathways: carotenoid biosynthesis (n=5 partitions; KEGG 00906), porphyrin and chlorophyll metabolism (n=4; KEGG 00860) and biosynthesis of secondary metabolites (n=5; KEGG 01110). These partitions supported gene swaps between lineages A3 and A5 (occurrence in 33 PEG trees), between lineages A7 and D2 (occurrence in 25 PEG trees) and between lineages C1 and C6 (occurrence in 126 PEG trees ; Figure 2A, Table 1, Table S6). The three partitions (A3-A5, C1-C6 and A7-D2) were consistently observed in 15 PEG trees, including 6 genes involved in carotenoid biosynthesis (*crtC*, *crtD*, *crtE*, *crtF*, *crtI, ppsR*), 7 genes in porphyrin and bacteriochlorophyll metabolism (*bchB*, *bchC*, *bchN*, *bch44.5kDa*, *pucC*, *pufH*, *pufM*) and two genes with unknown function (*rpc1322*, *rpc1324*; Table 1, Figure 2C). In addition, 9 genes involved in porphyrin and bacteriochlorophyll metabolism were observed in two partitions: A3-A5 and A7-D2 (*bchF*, *pufA* and *tspO*), A3-A5 and C1-C6 (*bchM*, *bchY* and *bchZ*) or A7-D2 and C1-C6 (*bchH*, *pufB* and *pufC*; Table 1, Figure 2C).

**Table 1:** List of 32 BCHL core PEGs. Genes were subdivided in subclusters BCHL1 and BCHL2 and ordered according to their position in CBMB20 genome. Following information is displayed for each PEG: gene abbreviation, gene code for phylogenetic analyses, RAST annotation, HGT likelihood (RES), KEGGs pathways (01100: Metabolic pathways; 01110: Biosynthesis of secondary metabolites; 00860: Porphyrin and chlorophyll metabolism; 02020: Two-component system signaling pathway; 00900: Terpenoid backbone biosynthesis; 00906 : Carotenoid biosynthesis) and observation of alternative partitions supporting HGT between lineages A3 and A5, C1 and C6 or A7 and D2.

| **Gene** | **Code** | **RAST Annotation** | **RES** | **KEGG(s)** | **A3-A5** | **C1-C6** | **A7-D2** |
|---|---|---|---|---|---|---|---|
| *Cluster BCHL1* |  |  |  |  |  |  |  |
| *bchD\** | CG_798 | Protoporphyrin IX Mg-chelatase subunit D (EC 6.6.1.1) | 0.028 | 01100; 01110; 00860 | na | na | na |
| *bchI* | CG_800 | Protoporphyrin IX Mg-chelatase subunit I (EC 6.6.1.1) | -0.008 | 01100; 01110; 00860 | - | X | - |
| *RPC_1324* | CG_1084 | Uncharacterized protein RPC_1324 | 0.077 | na | X | X | X |
| *acsF* | CG_599 | Mg protoporphyrin IX monomethyl ester oxidative cyclase (aerobic) (EC 1.14.13.81) | 0.045 | 01100; 01110; 00860 | - | X | - |
| *RPC_1322* | CG_1083 | Uncharacterized protein RPC_1322 | 0.025 | na | X | X | X |
| *pufH* | CG_744 | Photosynthetic reaction center H subunit | 0.017 | 02020 | X | X | X |
| *pucC* | CG_804 | PucC protein | 0.050 | na | X | X | X |
| *bchM* | CG_600 | Mg-protoporphyrin O-methyltransferase (EC 2.1.1.11) | 0.047 | 01100; 01110; 00860 | X | X | - |
| *bchL* | CG_514 | Light-independent protochlorophyllide reductase iron-sulfur ATP-binding protein ChlL (EC 1.3.7.7) | 0.059 | 01100; 01110; 00860 | - | X | - |
| *bchH* | CG_799 | Protoporphyrin IX Mg-chelatase subunit H (EC 6.6.1.1) | 0.068 | 01100; 01110; 00860 | - | X | X |
| *bchB* | CG_515 | Light-independent protochlorophyllide reductase subunit B (EC 1.3.7.7) | 0.037 | 01100; 01110; 00860 | X | X | X |
| *bchN* | CG_516 | Light-independent protochlorophyllide reductase subunit N (EC 1.3.7.7) | 0.060 | 01100; 01110; 00860 | X | X | X |
| *bchF* | CG_24 | 2-vinyl bacteriochlorophyllide hydratase BchF (EC 4.2.1.-) | 0.023 | 01110; 00860 | X | - | X |
| *ppsR* | CG_840 | Regulator of carotenoid biosynthesis; Transcriptional regulator, PpsR | 0.042 | na | X | X | X |
| *bchG* | CG_223 | Chlorophyll a synthase ChlG (EC 2.5.1.62) | 0.060 | 01100; 01110; 00860 | - | X | - |
| *bch44.5kDa* | CG_166 | Bacteriochlorophyll synthase 44.5 kDa chain | 0.038 | na | X | X | X |
| *bchP\** | CG_400 | Geranylgeranyl diphosphate reductase (EC 1.3.1.83) | 0.030 | 01100; 01110; 00860; 00900 | na | na | na |
| *tspO* | CG_1041 | Tryptophan-rich sensory protein precursor | 0.019 | 04080_04214 | X | - | X |
| *crtI* | CG_747 | Phytoene desaturase (neurosporene-forming) (EC 1.3.99.28) | 0.043 | 01100; 01110; 00906 | X | X | X |
| *Cluster BCHL2* |  |  |  |  |  |  |  |
| *pufC* | CG_743 | Photosynthetic reaction center cytochrome c subunit | 0.040 | na | - | X | X |
| *pufM* | CG_746 | Photosynthetic reaction center M subunit | 0.015 | 02020 | X | X | X |
| *pufL* | CG_745 | Photosynthetic reaction center L subunit | 0.020 | 02020 | - | - | - |
| *pufA* | CG_512 | Light-harvesting LHI, alpha subunit | 0.022 | 02020 | X | - | X |
| *pufB* | CG_513 | Light-harvesting LHI, beta subunit | 0.012 | 02020 | - | X | X |
| *bchZ* | CG_226 | Chlorophyllide reductase subunit BchZ (EC 1.18.-.-) | 0.048 | 01110; 00860 | X | X | - |
| *bchY* | CG_225 | Chlorophyllide reductase subunit BchY (EC 1.18.-.-) | 0.039 | 01110; 00860 | X | X | - |
| *bchX* | CG_224 | Chlorophyllide reductase subunit BchX (EC 1.18.-.-) | 0.035 | 01110; 00860 | - | - | - |
| *bchC* | CG_16 | 2-desacetyl-2-hydroxyethyl bacteriochlorophyllide A dehydrogenase BchC | 0.059 | 01110; 00860 | X | X | X |
| *crtF* | CG_478 | Hydroxyneurosporene methyltransferase (EC 2.1.1.210) | 0.071 | 01100; 01110; 00906 | X | X | X |
| *crtE* | CG_401 | Geranylgeranyl diphosphate synthase (EC 2.5.1.29) | 0.015 | 01100_01110_00900 | X | X | X |
| *crtD* | CG_591 | Methoxyneurosporene dehydrogenase (EC 1.14.99.-) | 0.041 | 01100; 01110; 00906 | X | X | X |
| *crtC* | CG_88 | Acyclic carotenoid 1,2-hydratase (EC 4.2.1.131) | 0.084 | 01100; 01110; 00906 | X | X | X |

\*Because at least two lineages were missing for *bchP* and *bchD* original trees, partition data for these genes were not included in this table

### HGT likelihood estimation based on modified RF distance (RES) and KEGG pathway enrichment

To quantify the degree of phylogenetic deviation of each PEG tree from the *Methylobacterium* consensus tree, we compared individual core PEG phylogenies with the consensus lineage tree using Robinson–Foulds (RF) distances (Figure S7). Raw RF distances were significantly associated with investigated properties of individual gene alignments and phylogenetic trees, including median nodal support (*Mns*; p<0.001), number of taxa per tree (*nTIP*; p<0.001), dN/dS ratio (*dNdS*; p<0.001), and number of polymorphic sites (*nPOL*; p<0.05; ANOVA, Table S7).

In the initial linear model, *Mns* explained most of the variance in RF distances (73.3%), followed by *nTIP* (6.9%), while *dNdS* and *nPOL* individually accounted for smaller proportions of variance (<0.3% each). Several interaction terms were also significant, including *dNdS*:*nPOL* and *dNdS*:*nPOL*:*TIP* (Table S7). Iterative model simplification retained all four main factors (*Mns*, *nTIP*, *dNdS* and *nPOL*) together with significant interaction terms in the final model, all with similar explanation of RF variance (Table S7). Residual RF values (RES) of the simplest model were no longer associated with *dNdS*, *nPOL*, *Mns* or *nTIP* (Figure S7) and were subsequently used as a proxy for HGT likelihood in downstream analyses.

We compared average RES values among KEGG pathways to identify signatures of phylogenetic incongruences across functional pathways (Figure S8; Table S5). Among the 134 KEGG pathways identified across core PEGs, two showed significantly higher RES values than expected by chance (1.10^6^ permutations tests, BH correction for multiple testing): carotenoid biosynthesis (n=6 PEGs; KEGG 00906; mean RES = 0.042; adjusted p=0.024) and porphyrin metabolism (n=46 PEGs, KEGG 00860; mean RES = 0.013; adjusted p=0.046; Table S5).

Together, our analysis so far indicates that genes involved in bacteriochlorophyll and carotenoid metabolism are more likely to have been horizontally transferred, resulting in incongruent phylogenetic inferences with other *Methylobacterium* core genes.

### Gene cluster enrichment in HGT (based on RES)

To identify gene clusters potentially enriched in horizontal gene transfer (HGT), we aggregated genes into clusters according to different CCL values (in the range 8-24 lineages) and compared their average RES values to null expectations generated by permutations (Table S8; Figure 3). Two conserved clusters displayed significantly elevated RES values, indicating unusually strong phylogenetic incongruence and therefore increased HGT likelihood (Figure 3; 1.10^6^ permutations; adjusted p<0.001). One large cluster included 32 PEGs involved in carotenoid and bacteriochlorophyll biosynthesis, (thereafter BCHL cluster; Table 1; Table S8). The BCHL cluster formed one block for CCL values in the range 8-13 and was subdivided in two conserved sub-clusters for CCL values in the range 14-24. The BCHL1 cluster contained most components for bacteriochlorophyll *a* biosynthesis (19 genes: *acsF, bch44.5kDa, bchB, bchD, bchF, bchG, bchH, bchI, bchL, bchM, bchN, bchP, crtI, ppsR, pucC, pufH, RPC_1322, RPC_1324, tspO*; Table 1 ;Figure S9). The BCHL2 cluster contained most genes for carotenoid biosynthesis, the type-II reaction center, and light harvesting complex of the photosystem (13 genes: *bchC, bchX, bchY, bchZ, crtC, crtD, crtE, crtF, pufA, pufB, pufC, pufL, pufM*; Table 1 ;Figure S9). Separately, BCHL1 and BCHL2 were also enriched in HGT likelihood (mean RES = 0.040 ± 0.021 and 0.038 ± 0.023, respectively; adjusted p<0.01). The second cluster corresponded to four PEGs associated with osmoregulation (thereafter KDP cluster; Table S8), including the osmosensitive sensor kinase gene *kdpD* and the potassium-transporting ATPase genes *kdpA*, *kdpB* and *kdpC* (mean RES = 0.088 ± 0.020; adjusted p<0.001; Figure 3, Table S8) and was conserved across all *Methylobacterium* lineages (CCL range 8-24).

**Figure 3:**
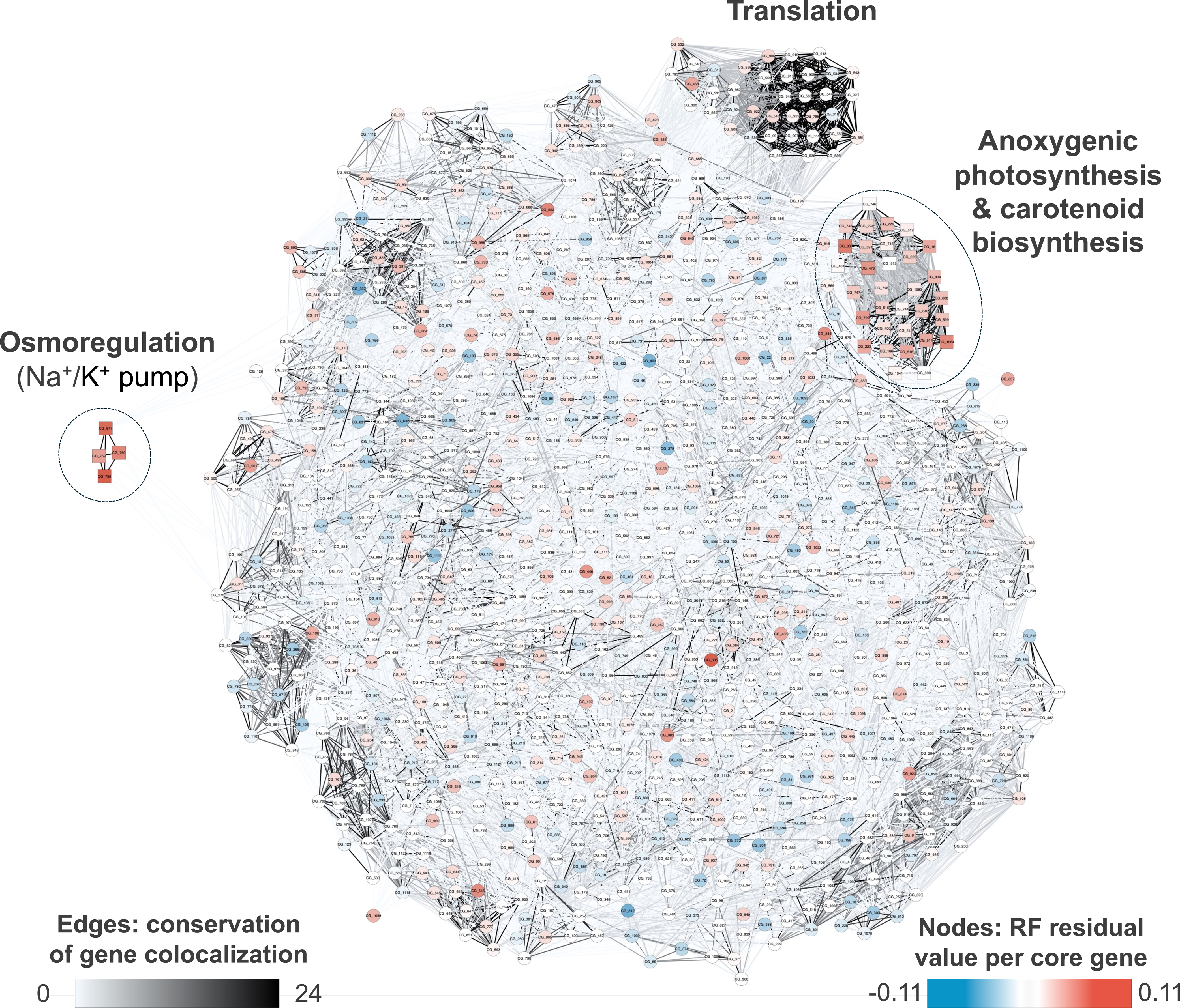
Phylogenetic discordance is concentrated in conserved functional gene clusters. The colocalization network of core genes in *Methylobacterium* genomes (cytoscape) shows that gene clusters involved in osmoregulation (sodium/potassium pump), photosynthesis (bacteriochlorophyll), and UV protection (carotenoids) are conserved in *Methylobacterium* while being enriched in HGT. Genes are represented by circles colored according to residual RF distance (red indicate high HGT likelihood; square indicate genes for which phylogeny was redone, including related Hyphomicrobiales in order to identify potential inter-genera HGT). Edge thickness and color indicate the proportion of genomes (weighted by the number of genomes within species and by the number of species within lineages) where the two genes colocalized (i.e. were found within the same 25 kb region). Doted frames indicate cluster of genes (conserved colocalization among a majority of *Methylobacterium* lineages) that have higher residual RF value than expected by chance (p < 0.001; 100,000 permutations, BH correction).

### Detection of HGT using nucleotide polymorphism patterns (BAPS)

To complement phylogenetic discordance analyses, we investigated nucleotide polymorphism patterns across the 1,099 core PEGs using hierarchical Bayesian Analysis of Population Structure (BAPS). BAPS analyses were performed independently for each gene alignment, assuming 24 populations, corresponding to the number of *Methylobacterium* lineages (Figure S10A-B).

For each of the 1,099 core PEGs, BAPS lineages were compared with both the consensus ASTER phylogeny and the corresponding gene tree and classified as congruent with vertical inheritance (BAPS lineages congruent with both the species and the gene tree), indicative of putative HGT (incongruence with the species tree), or undetermined (incongruence between BAPS lineages and the gene tree; Figure 4, Figure S10C). Undetermined cases were more frequently observed for BAPS lineages containing deep-branching *Methylobacterium* lineages (D5; D6; B3; B4; A3; A4; A6; A7; Figure S10C; Figure 4D).

**Figure 4:**
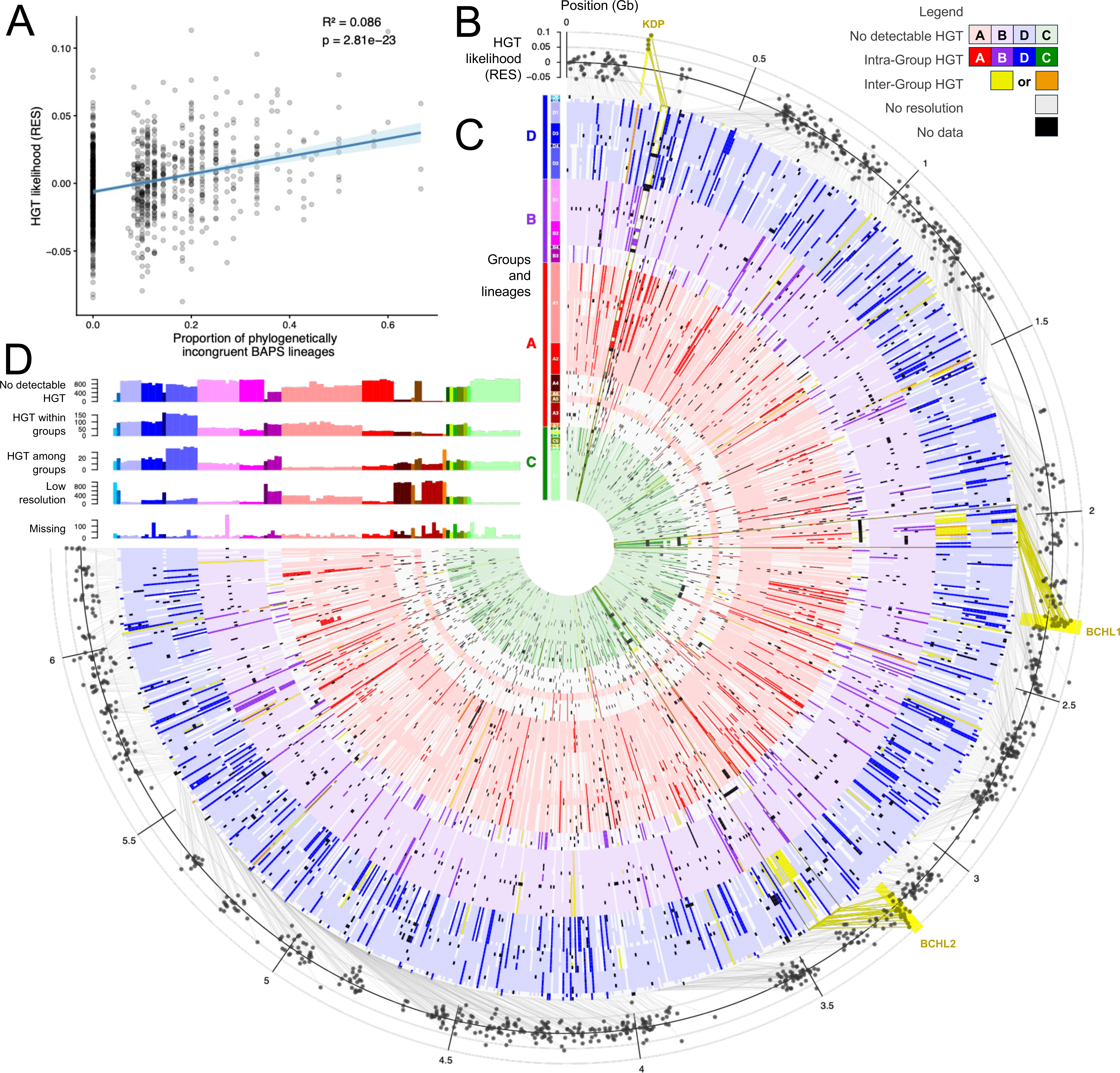
Bayesian-based assignment of *Methylobacterium* core PEGs ancestry based on nucleotide polymorphism is congruent with gene trees. A) The proportion of lineages inferred by BAPS that are incongruent with the consensus ASTER *Methylobacterium* tree is significantly correlated with HGT likelihood (RES) per PEG tree (linear model, F-test). B) HGT likelihood per core PEG distribution along the CBMB20 reference genome. Gene clusters that have higher residual RF values than expected by chance and correspond to BCHL and KDP clusters (see Figure 3) are highlighted in yellow. C) BAPS assignment of *Methylobacterium* species (in ordered rows according to the ASTER species tree) in phylogroups for each core PEG (columns, ordered to their position along the CBMB20 genome, grey links). Colors and shades indicate congruence between BAPS analysis, core PEG tree topology and the consensus tree phylogeny (legend on top right). Putative HGT among phylogroups are enriched in BCHL and KDP clusters. D) Absolute number of core PEGs per congruence category between BAPS analysis and per core PEG congruence with the ASTER consensus tree.

Overall, we detected putative HGT in 585 out of 1,099 PEGs, of which 495 involved exchanges among lineages within phylogroups exclusively, 33 among phylogroups exclusively, and 57 PEGs with putative HGT both among and within phylogroups (Figure S10D). Putative HGT within phylogroups mostly involved lineages D4 (observed in 165 PEGs), D2 (151 ± 6), D3 (97 ± 5), D1 (96 ± 2), D5 (90), B4 (89) and/or B3 (78 ± 2; Figure S10C; Figure 4D). HGT among phylogroups mostly involved lineages D2 (in 37 ± 2 PEGs), B3 (20 ± 4) and/or A7 (34; Figure S10C; Figure 4D). Overall, classifications inferred from BAPS were broadly consistent with HGT likelihood estimated from corrected RF distances per core PEG (RES), with the proportion of BAPS lineages classified as putative HGT (intra- and inter-phylogroup combined) significantly correlated with RES (linear model: R² = 0.082, p <<0.001; Figure 4A).

When mapped along the CBMB20 reference genome, BAPS classifications revealed substantial variation among genomic regions (Figure 4B-C). Hotspots of PEGs with elevated proportions of incongruent lineage assignments corresponded to BCHL and KDP clusters (Figure 3; Figure 4B-C). Across the BCHL clusters, 30 out of 32 PEGs showed evidence of intra-phylogroup and/or inter phylogroup HGT according to BAPS classification, whereas 4 out of 4 PEGs from the KDP cluster showed similar patterns (Figure 4C, Figure S10D).

### HGT in BCLH clusters in Methylobacterium and related Hyphomicrobiales

To characterize the evolutionary history of the bacteriochlorophyll and carotenoid biosynthesis cluster (BCHL) identified as strongly enriched in phylogenetic incongruence and HGT signals, we extended comparative analyses beyond *Methylobacterium* to closely related Hyphomicrobiales lineages inhabiting the phyllosphere and related environments. We re-analyzed the 32 BCHL-associated core genes (Table 1) across the 293 *Methylobacterium* genomes together with 167 genomes and metagenome-assembled genomes (MAGs) from related Hyphomicrobiales families and genera (Table S9).

BCHL clusters were broadly distributed across distantly related Hyphomicrobiales, although their occurrence was highly uneven among genera and did not follow the consensus phylogeny (Figure 5A-B). Within *Methylobacterium*, BCHL genes were detected in 113 out of 116 species (Figure 5B; Figure S11). Outside *Methylobacterium*, BCHL clusters were mostly observed in dominant phyllosphere-associated Hyphomicrobiales (Leducq et al. 2026), including the undescribed *RH_AL1* group (23 out of 23 genomes) and *Lichenibacterium* (13 out of 18; Figure 5A-B; Table S9).

**Figure 5:**
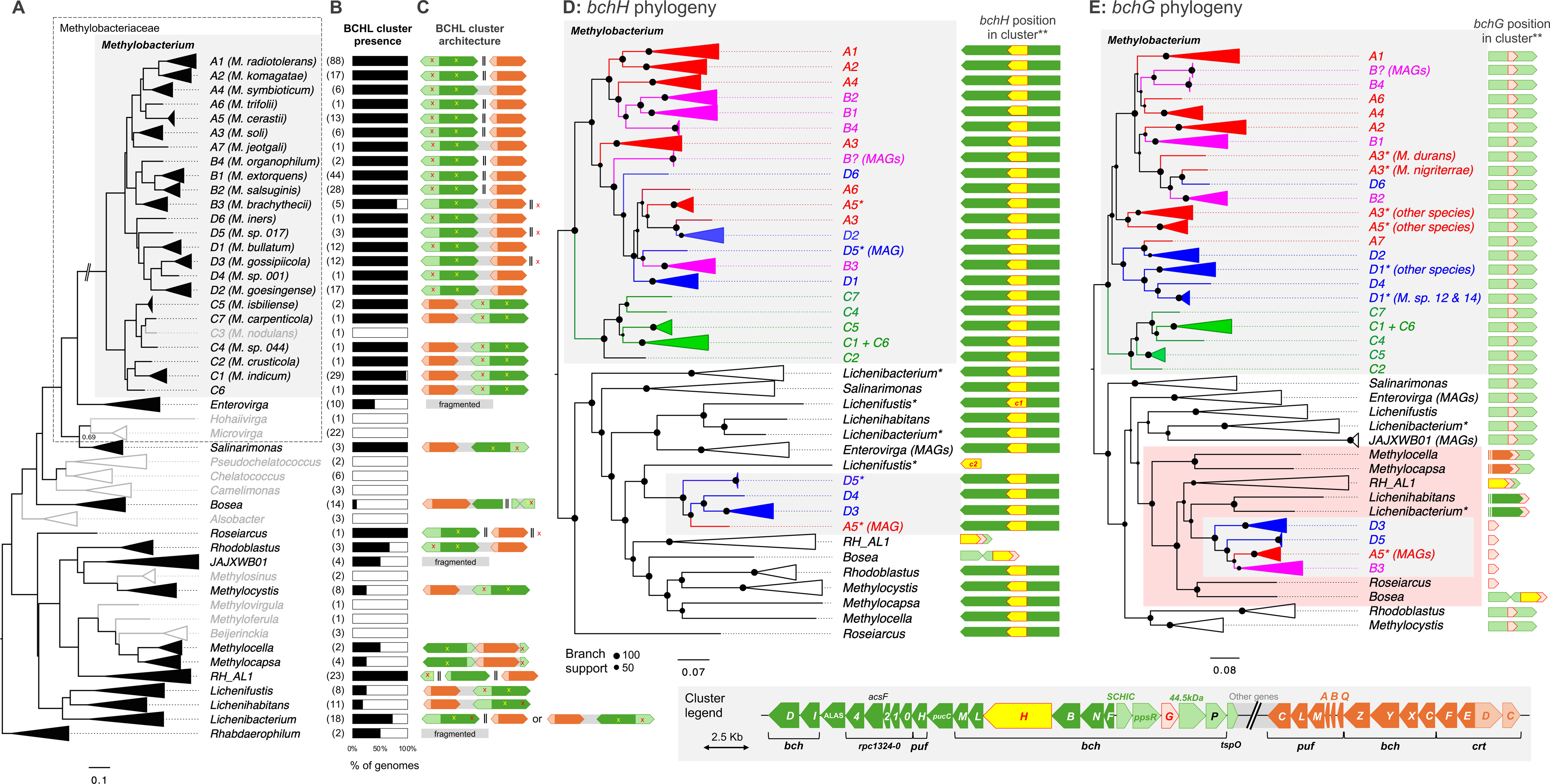
Bacteriochlorophyll and carotenoid gene cluster evolution driven by HGT and dynamic cluster architecture in *Methylobacterium* and related genera. A) Consensus Hyphomycrobiales phylogeny combining this study (*Methylobacterium* lineages) and our previous work on related Hyphomicrobiales genera (Leducq et al. 2026). All nodes have branch support = 1 (local posterior probability) unless specified. For each clade, the total number of genomes that were considered for BCHL cluster analysis is indicated in parentheses. B) For each *Methylobacterium* lineage and related Hyphomicrobiales genera, proportion of genomes with a BCHL gene cluster. C) BCHL gene cluster architecture (simplified from Clinker) comparison among *Methylobacterium* lineages and related Hyphomicrobiales genera (detailed architecture in legend). Colored arrows indicate well-conserved subclusters of genes across genera. Grey frames indicate that subclusters are fused in a single cluster (separated by several accessory genes often related to BCHL). Double lines indicated that genes are localized in different regions of the genome. Red and yellow marks indicate *bchG* and *bchH* gene positions, respectively. Alternative BCHL architectures are displayed for *Lichenibacterium*. D-E) *bchH* and *bchG* gene phylogenies and gene position in the cluster. Asterisks indicate polyphyletic lineages and genera. For *bchG*, phylogenetic incongruence caused by a monophyletic group (red frame) correlated with the unusual gene localisation for genomes within this group.

The fusion between clusters BCHL1 and BCHL2 was variable across *Methylobacterium* species and lineages (Figures S11). BCHL1 and BCHL2 were separated in most species of phylogroups A and B but were fused in a single cluster for deep-branching lineages A4, A7 and B3. BCHL1 and BCHL2 were also fused in most species of phylogroups C and D, but separation was unevenly observed for some species. The BCHL12 fusion in representative genomes of each *Methylobacterium* lineage could be confirmed with clinker analyses (Figure S12; Figure 5C). BCHL1 cluster organisation was conserved between genes *bchD* and *tspO*, with few lineage-specific variations. The *bchG* gene was located outside the BCHL1 cluster exclusively in lineages B3, D5 and D3. Genes *bchD* and *bchI* were located outside the BCHL1 cluster exclusively in lineages B1 and B2 (in those cases, BCHL1 ended with the gene *ALAS*). BCHL2 cluster organisation was conserved and consistently flanked by genes *pufC* and *crtC* across all *Methylobacterium* lineages. The fusion between clusters BCHL1 and 2 observed in some lineages of phylogroups A, B, and D followed the same pattern, with BCHL1 and BCHL2 connected at their *bchD*/*ALAS* and *crtC* ends, respectively, through the addition of genes also involved in carotenoid biosynthesis (*crtB*, *crtI*, *ABH*). On the other hand, in phylogroup C, BCHL1 and 2 were inverted, with the fusion occurring between *tspO* and *pufC* ends, through the uneven addition of genes (1-4 depending on the lineage, including *c-553*).

We observed substantial variation in BCHL cluster architecture among Hyphomicrobiales genera and within *Lichenibacterium* (Figure 5C, Figure S13). Fusions between clusters BCHL1 and 2 were observed within all genera except *Roseiarcus* and *RH_AL1*. All possible fusion patterns between clusters BCHL1 and BCHL2 could be observed. A pattern typical of *Methylobacterium* phylogroups ABD was observed in *Rhodoblastus*, while the pattern typical of *Methylobacterium* phylogroup C was observed in *Methylocystis*, *Lichenifustis,* and *Lichenihabitans*. Alternative fusion patterns were observed in *Salinarimonas* and *Lichenibacterium*. BCHL1 and 2 substructure was overall similar to that observed in *Methylobacterium*, with few variations. Differences among genera were notably marked by extensive variations in the position of *bchG* and surrounding genes in the cluster BCHL1 (Figure 5C).

To identify putative HGT events between *Methylobacterium* and related Hyphomicrobiales genera, we reconstructed phylogenies for each BCHL gene individually, including homologs from related Hyphomicrobiales genomes. In 27 out of the 32 BCHL gene phylogenies, *Methylobacterium* formed a monophyletic group. We further examined phylogenies of genes where *Methylobacterium* was not monophyletic: *bchH*, *bchG*, *pufC*, *pufM* and *pufL*. Gene trees of *bchH* (protoporphyrin IX Mg-chelatase subunit H) and *bchG* (chlorophyll a synthase *ChlG*) displayed strong topological discordance relative to the consensus Hyphomicrobiales phylogeny, with reshuffled phylogenetic relationships among genera. *Methylobacterium* and *Lichenibacterium* consisted of several clades in both cases (Figure 5D-E). Both genes were found in a single copy in all genomes with BCHL clusters, except for *Lichenifustis*. In *Lichenifustis*, the *bchH* gene appeared duplicated and the extra copy was located outside the BCHL cluster. For *bchH*, gene copies from lineages A5, D3, D4 and A5 formed a monophyletic group outside the core *Methylobacterium* clade, branching with the *Lichenifustis* extra copy (nodal support =99; Figure 5E). For *bchG*, gene copies from lineages D3, D5, B3 and A5 formed a monophyletic group outside the core *Methylobacterium* clade, branching with *Lichenibacterium*, *Lichenihabitans*, *RH*_*AL1, Bosea and Roseiarcus* (nodal support =99; Figure 5E). In both gene phylogenies, the core *Methylobacterium* clade appeared rooted on phylogroup C, which remained monophyletic; while phylogenetic relationships were completely reshuffled among and within phylogroups A, B and D.

In gene trees of *pufC* (photosynthetic reaction center cytochrome *c* subunit), *pufM* (photosynthetic reaction center M subunit) and *pufL* (photosynthetic reaction center L subunit), *Methylobacterium*, *RH_AL1* and *Lichenibacterium* consisted of several clades (Figure S14). In *Methylobacterium*, this pattern consistently resulted from the duplication of the *pufC-pufM-pufL* gene tandem observed in some genomes of lineages A1 (5 out of 88 genomes) and A2 (4 out of 17 genomes), and a duplication of *pufC* solely observed in lineage A3 (3 out of 6 genomes). In *RH_AL1*, we observed a similar pattern of complete or partial duplication of the *pufC-pufM-pufL* gene tandem in 14 out of 23 genomes. In the three gene phylogenies, *RH_AL1* copies were distributed in two monophyletic clades (hereafter clades *pufC2* and *pufC3*, *pufL2* and *pufL3*, *pufM2* and *pufM3*). Each *Methylobacterium* genome with the observed duplication possessed at least one copy with congruent phylogenetic placement in its native lineage within the core *Methylobacterium* clade (hereafter clade *pufC1*, *pufL1* and *pufM1*), while the extra copies of lineages A1 and A2 systematically formed a monophyletic clade with *RH_AL1* copies *pufC2*, *pufL2* and *pufM2* (nodal support =100) and the extra *pufC* copy of lineage A3 formed a monophyletic clade with *RH_AL1* copy *pufC3* (nodal support =67). We observed *pufL* and *pufM* in a majority of *Lichenibacterium* genomes (11 out of 18). Most copies (10 out of 11) from these genomes belonged to *pufL2* and *pufM2 clades* (nodal support =100) and lacked *pufC*. In the only *Lichenibacterium* genome with a complete *pufC-pufM-pufL* tandem, gene copies formed a monophyletic group with copies from a few *Lichenihabitans* genomes (2 out of 11) and *Lichenifustis* genomes (2 out of 7) possessing a complete BCHL cluster (clades *pufC4*, *pufL4* and *pufM4/5*).

## Discussion

Our results reveal that the evolutionary history of the *Methylobacterium* core genome is far more heterogeneous than expected under a strictly vertical model of genome evolution. Rather than separating clearly into congruent versus incongruent gene sets, gene tree comparison, phylogenetic network, and nucleotide polymorphism pattern analyses consistently showed that most core genes displayed some degree of deviation from the consensus species phylogeny. First, phylogenetic network reconstruction indicated that conflicting phylogenetic signals are widespread across the core genome (Figure 2; Table S6). While the backbone of the *Methylobacterium* evolutionary network was congruent with the consensus *Methylobacterium* tree, with the 21 corresponding partitions observed in almost a majority (48.1 ± 23.9%) of core PEG trees, the root of the network was poorly resolved, with alternative possible relationships among phylogroups A, B, D and C observed almost equally but at low frequencies among core PEG trees (7.8-18.2%). This observation is congruent with our previous observations based on a smaller set of core genes (*n*=384) and with our hypothesis that early *Methylobacterium* history was obscured by HGT among phylogroups (Leducq, Sneddon, et al. 2022). Similarly, we consistently observed 369 alternative relationships among lineages, supporting alternative evolutionary scenarios among *Methylobacterium* lineages, within (3.5-8.8% of core PEG trees) and among (0.1-0.6%) phylogroups, indicating extensive gene swaps prior to and after *Methylobacterium* phylogroups and lineage divergence and radiation. The proportion of predicted gene swaps observed between lately diverging phylogroups A, B and D (54.8 %; range 35.0-70.8%) was significantly higher than proportions observed for exchanges with the most basal phylogroup C (9.8%; range 4.2-14.6%). This result confirms our previous observation that phylogroup A, B and D share a larger pool of genes than with phylogroup C, and further suggest the role of gene exchanges in blurring the phylogenetic resolution among the three groups (Leducq, Sneddon, et al. 2022). This lack of resolution and more broadly, to which extent phylogenetic discordances among genes are considered in *Methylobacterium* phylogenetic reconstruction models, may partially explained recent taxonomic confusions in the genus (Alleman et al. 2025). From and ecological perspective, the more frequent gene exchanges detected between phylogroups A, B and D might reflect their overlapping niche in the phyllosphere (Alleman et al. 2025), as ecological co-occurrence promote physical interactions among bacteria, including HGT (Dmitrijeva et al. 2024), while exposure to similar local conditions may impose similar selective pressures on horizontally acquired genes (Acar Kirit et al. 2022), favouring their retention across phylogroups. In contrast, the divergence of the basal phylogroup C likely reflects its adaptation to soil and rhizosphere environments and its ecological separation from other Methylobacterium lineages (Alleman et al. 2025). Second, we used an independent assessment of evolutionary discordance based on nucleotide polymorphism patterns, further supporting putative HGT signals in more than half of all core PEGs (53.2%; Figure 4, Figure S10). Although most of these signatures (50.2%) were detected among lineages within phylogroups, suggesting that gene exchange may be a recurrent process throughout the diversification of lineages, a significant fraction of genes (8.2%) also showed signal of exchanges among phylogroups. Third, using a modified Robinson-Fould (RF) distance metric accounting for core PEGs evolutionary characteristics and occurrence biases (Figure S7), we developed an original index quantifying core PEGs tree topology deviation from the *Methylobacterium* consensus lineage tree (Figure 3), confirming both network-based inference and BAPS analyses that core PEGs followed a continuum of alternative evolutionary histories. Although the correlation between BAPS-derived HGT estimates and corrected RF distances (RES) was modest (R² = 0.082), it was highly significant and was detected despite the fundamentally different assumptions underlying the two approaches. Whereas RF-based analyses rely on topological incongruence among phylogenetic trees, BAPS infers population structure directly from nucleotide polymorphism patterns. The convergence of these independent approaches suggests that phylogenetic discordance across the core genome is not solely driven by stochastic variation in phylogenetic reconstruction, but instead by alternative evolutionary signals among core PEGs, which is consistent with theoretical expectations that gene tree discordance is pervasive in genomic datasets (Maddison 1997; Degnan and Rosenberg 2009). While increasing the number of core PEGs statistically strengthens the genus consensus evolutionary tree reconstruction, our study combining complementary approaches highlights that core-genome-based phylogenetic proxies hide extensive variations in alternative core gene evolutionary histories, supporting a reticulate rather than purely vertical evolution of the core genome.

Although evolutionary heterogeneity extends across the entire core genome, only a small subset of PEGs displayed levels of discordance sufficiently elevated to emerge as significant outliers in different analyses. Gene clusters enriched in alternative network evolution (Figure 2), phylogenetic incongruence (Figure 3) and polymorphism patterns (Figure 4) were consistently and significantly associated with genes involved in bacteriochlorophyll and carotenoid biosynthesis pathways (Figure 2; Table S5). These functions are central to light harvesting and protection against oxidative and UV stress— two major constraints of the phyllosphere environment. The non-random association of these functions with phylogenetic incongruence contrasts with expectations under purely neutral processes such as incomplete lineage sorting and instead suggests evolutionary dynamics linked to ecological adaptation. Similar patterns have been reported previously in *Methylobacterium*, where high divergence among species in photosynthesis-related genes was interpreted as genomic innovation at the cluster level (Zervas et al. 2019). Our results rather suggest that this divergence reflects horizontal gene transfer and is consistent with observations in other anoxygenic photosynthetic Hyphomicrobiales like *Bradyrhizobium* and *Roseobacter*, and more broadly across bacteria, where phototrophic functions follow complex and often reticulate evolutionary trajectories shaped by horizontal gene transfer (Liu et al. 2019; Ward and Shih 2021; Avontuur et al. 2023). In addition to photosynthesis-related functions, we identify a second gene cluster (KDP) enriched in phylogenetic discordance and in BAPS analyses, comprising genes involved in potassium transport and osmoregulation (Figure 3; Figure 4; Table S8; (Rhoads et al. 1978)). In contrast to the BCHL cluster, the KDP cluster is highly conserved in both gene content and genomic organization across lineages yet exhibits elevated phylogenetic incongruence. Potassium transport systems are critical for maintaining osmotic balance under rapid environmental changes (Sleator and Hill 2002). In the phyllosphere, where microorganisms experience repeated cycles of hydration and desiccation, such functions are likely critical for survival. The enrichment of phylogenetic discordance in both the BCHL and KDP clusters might reflect adaptation to light exposure and osmotic stress, two key constraints affecting bacterial growth and survival in the phyllosphere.

We show that phylogenetic similarity among genes decreases with genomic distance (i.e. physical separation in the genome), with genes that co-localize across genomes tending to display more similar evolutionary histories (Figure S5). Functional similarity (based on KEGG pathways) follows a similar pattern, indicating that physically linked genes are also more likely to be functionally related. These observations support the view that bacterial genomes are structured into functionally cohesive units whose evolutionary trajectories may be partially distinct from the genomic background (Rocha 2008; Wiedenbeck and Cohan 2011). The strong enrichment of HGT signals at the cluster level (Figure 3; Table S8), particularly for the BCHL and KDP clusters, is consistent with the idea that such gene sets may coevolve and be horizontally transferred as integrated modules (Dilthey and Lercher 2015). Those enrichments also indicate that, although phylogenetic discordance is widespread across the core genome, the signature is stronger in specific functional categories associated with adaptation to life in the phyllosphere. In this context, BCHL and KDP should be viewed as extreme examples of a broader continuum of evolutionary dynamics, where horizontal gene and operon transfers are frequent but heavily filtered by purifying and positive selection (Power et al. 2021; Nguyen et al. 2022).

The BCHL cluster provides a particularly illustrative example of these dynamics. Across *Methylobacterium* lineages, bacteriochlorophyll and carotenoid biosynthesis genes are organized into conserved subclusters (BCHL1 and BCHL2) that exhibit variable architectures, including lineage-specific fusion and separation patterns (Figure 5C; Figures S11–S13). Despite extensive genome rearrangements across the genus, the internal organization of these clusters remains highly conserved, suggesting functional constraints. This is consistent with the functions of genes in BCHL1, mostly involved in bacteriochlorophyll biosynthesis, while BCHL2 genes are predominantly involved in carotenoid biosynthesis and light-harvesting complexes (Figure S9). At the same time, the BCHL cluster displayed substantial rearrangements and phylogenetic incongruences, both within *Methylobacterium* and across related Hyphomicrobiales (Figure 5E–F), consistent with repeated and potentially modular horizontal exchanges. The highly dynamic evolution of the BCHL cluster in terms of structure and gene exchange has recently been highlighted in *Bradyrhizobium*, another plant-associated Hyphomicrobiales genus (Avontuur et al. 2023), supporting the role of HGT as a common source of innovation in anoxygenic photosynthesis in Hyphomicrobiales. While anoxygenic photosynthesis is not ancestral and was probably acquired multiple times in *Bradyrhizobium* (Avontuur et al. 2023), our results suggest a different scenario for *Methylobacterium*. The detection of the BCHL gene cluster in most of the analyzed genomes (290/293; Figure 5A–B; Table S9) suggests that anoxygenic photosynthesis is an ancestral feature in *Methylobacterium*. Phylogenetic analysis extended to related Hyphomicrobiales showed that *Methylobacterium* was monophyletic for a majority (27 out of 32) of BCHL genes, further supporting that the BCHL cluster was acquired only once during *Methylobacterium* evolution, followed by repeated and punctuated gene exchanges between lineages as attested by extensive phylogenetic incongruences observed within the genus.

The distribution of BCHL clusters across Hyphomicrobiales is patchy and does not follow the consensus phylogeny (Figure 5A–B; Table S9), with some genera systematically harboring these pathways while others lack them entirely. This pattern is consistent with the phylogenetically discontinuous distribution of anoxygenic photosynthesis across bacteria (Ward and Shih 2021), a trend also suggested within Hyphomicrobiales (Zervas et al. 2019). Our results extend this observation to Hyphomicrobiales closely related to *Methylobacterium*, complementing previous work in *Bradyrhizobium* (Avontuur et al. 2023) showing repeated independent acquisitions of photosynthesis gene clusters. Moreover, similarities in cluster architecture across distantly related taxa, combined with phylogenetic incongruence at the gene level, are consistent with recurrent exchanges and functional constraints across lineages inhabiting related environments. These observations support the idea that phototrophic functions may form part of a shared functional repertoire that is intermittently exchanged among phylogenetically distinct but ecologically overlapping bacteria (Ward and Shih 2021).

Further resolution of these patterns at the gene level highlights particularly strong discordance for *bchH* (Figure 5D) and *bchG* (Figure 5E), both within *Methylobacterium* and across Hyphomicrobiales. These genes encode key enzymatic steps in the bacteriochlorophyll biosynthesis pathway, corresponding to early (*bchH*; Mg-chelatase subunit) and late (*bchG*; chlorophyll synthase) stages of pigment formation (Figure S9) (Proctor et al. 2022). Incongruence in the phylogenetic signal of *bchG* is associated with shifts in its genomic context. In *Methylobacterium* lineages displaying discordant *bchG* phylogenies, the gene is systematically located outside its conserved position within the BCHL1 cluster or in rearranged cluster configurations. This association between phylogenetic incongruence and gene position is consistent with the proposed role of HGT in BCHL cluster reorganization reported at a larger evolutionary scale in photosynthetic bacteria (Ward and Shih 2021), and might reflect the impact of HGT on reshuffling the organization of BCHL genes. We observed a similar pattern for genes encoding core transmembrane protein subunits of the photosynthetic reaction center (*pufC*, *pufL,* and *pufM*; Figure S14), typical of purple bacteria (Imhoff et al. 2018). In contrast to *bchH* and *bchG*, phylogenetic incongruence in these genes is associated with the presence of multiple gene copies, resulting from lineage-specific duplications. In *Methylobacterium*, duplicated copies consistently include one lineage-congruent copy and one alternative copy clustering with homologs from other leaf-associated Hyphomicrobiales. These additional copies form well-supported monophyletic groups across distinct taxa, suggesting repeated exchange of specific gene variants rather than complete cluster transfer.

Our results indicate that photosynthetic gene evolution in these lineages involves not only large-scale cluster dynamics but also gene-specific duplication and exchange. Within Hyphomicrobiales, phylogenetic discordance, duplication, and position in clusters observed for BCHL genes *bchG*, *bchH*, *pufC*, *pufM,* and *pufL* are particularly marked among taxa that frequently co-occur in the phyllosphere. *Methylobacterium* species, Lichenihabitantaceae, and *RH_AL1* are abundant and frequently co-occur at fine spatial scales on plant surfaces (Knief et al. 2010; Leducq, Seyer-Lamontagne, et al. 2022; Leducq et al. 2026). Such ecological overlap is expected to facilitate genetic interactions and may increase opportunities for gene exchange, as described in other microbial systems (Polz et al. 2013; Woods et al. 2020). In the context of *Methylobacterium* in the phyllosphere, horizontal gene transfer may contribute to the maintenance or reshaping of adaptive traits, enabling bacteria to respond to fluctuating environmental conditions such as UV exposure, desiccation, and nutrient limitation.

Additional support for dynamic evolutionary boundaries within the genus comes from patterns of genomic divergence. Lineage delimitation based on ANI does not reveal a single consistent threshold separating evolutionary units, with lineage boundaries typically falling within a range of 84–95% identity (Figure 1B; Figure S3), as previously reported (Alleman et al. 2025). This absence of a clear-cut boundary is consistent with the idea that evolutionary divergence within *Methylobacterium* may not follow a strictly clonal trajectory but instead reflects a more complex history potentially involving gene flow among partially differentiated lineages.

While our results are consistent with horizontal gene transfer as a major contributing factor of phylogenetic discordance, alternative explanations must also be considered. Phylogenetic heterogeneity can arise from incomplete lineage sorting, variation in evolutionary rates across genes, or limitations in phylogenetic inference. However, several observations argue against purely stochastic explanations. First, discordance is structured by function rather than being randomly distributed. Second, genes within conserved clusters tend to share similar phylogenetic histories. Third, independent analytical approaches—including gene tree discordance, phylogenetic network reconstruction, and cluster-based enrichment—converge on similar patterns. Together, these results support a scenario in which horizontal gene transfer contributes to shaping the evolution of specific functional categories within the core genome, while remaining conservative in the absence of direct inference of transfer events. An additional limitation relates to the definition of the core genome itself. To accommodate the large number of draft assemblies included in this study, core genes were defined using deliberately relaxed presence thresholds designed to minimize artefactual gene absences caused by assembly fragmentation rather than biological variation. Although this strategy did not affect the detection of phylogenetic discordance itself, and even increased the robustness of comparative analyses, it also resulted in the inclusion of a small number of genes that may be more appropriately described as components of a relaxed-core genome. The BCHL cluster illustrates this point: being absent from only three out of 116 *Methylobacterium* species, it is identified as part of the genus core genome under our criteria, but is notably absent in some related genera.

Finally, further work should clarify mechanisms underpinning gene exchange and diversification of anoxygenic photosynthesis in the phyllosphere. Previous work in anoxygenic photosynthetic marine bacteria like *Roseobacter* and *Jannaschia* (Alphaproteobacteria, Rhodobacterales) suggested that HGT and recombination in BCHL clusters occurred through mobile genetic elements that disseminate and recombine modular gene clusters across lineages (Liu et al. 2019).

Our results suggest that core genomes—traditionally viewed as reflecting a largely vertical evolutionary history—may exhibit substantial phylogenetic heterogeneity. In *Methylobacterium*, this heterogeneity is concentrated in genes associated with ecological adaptation to the phyllosphere, indicating that selective pressures acting in specific environments may promote more dynamic evolutionary trajectories for these functions. More broadly, these results support a model in which bacterial genomes evolve as mosaics of genes with distinct evolutionary histories, shaped by the interplay between vertical inheritance, ecological selection, and genetic exchange.

## Conclusion

This study provides evidence that the diversification of key metabolic functions in *Methylobacterium* is consistent with a combination of vertical inheritance and dynamic gene exchange. The enrichment of phylogenetic discordance within clusters involved in photosynthesis and stress tolerance suggests that these functions have been repeatedly reshaped during *Methylobacterium* diversification, potentially contributing to adaptation to life on plant leaves. More generally, our findings highlight the importance of considering phylogenetic heterogeneity within genomes to better understand the processes generating functional diversity in microbial communities.

## Materials and Methods

All analyses were conducted in R (Team R Developement Core 2005), unless specified.

### Rice-associated Methylobacterium genome assembly

We performed *de novo* assembly for 69 putatively axenic *Methylobacterium* isolates from India (Sanjenbam et al. 2020)(Table S2). This collection included strains isolated in 2016 and 2017 from rice plant (*Oryza sativa*) phyllosphere (n=65; stems, seeds and leaves), from field soil (n=2) and from grass growing adjacent to rice plants (n=2) (Table S2). Illumina Hiseq paired-end 100bp short-read libraries were produced in 2018 (SR18) for all isolates; seven isolates were re-sequenced in 2020 (SR20). Short-read assemblies were generated with MEGAHIT v1.2.9 (Li et al. 2015) (default parameters except m = 0.5 and t = 10) to improve resolution when multiple related genomes could be present in a single assembly. Whenever available, three assemblies per isolate were generated — SR18 alone, SR20 alone, and a combined SR18+SR20 assembly — to assess library compatibility and detect potential cross-library contamination. In four isolates (MANGS3, APS3.4, KUML1, PML2), SR18 and SR20 libraries corresponded to distinct genomes and were therefore assembled separately, yielding a total of 73 assemblies.

The 73 assemblies were screened for contamination. In 11 of them, the genome exceeded the expected *Methylobacterium* size range (4.3–7.8 Mb; (Leducq, Sneddon, et al. 2022)) or displayed multimodal coverage and/or GC distributions, consistent with the presence of mixed genomes (Figure S1). These assemblies were further partitioned by reassigning scaffolds to putative genomes based on scaffold coverage and/or GC content, generating 2-3 unimodal assemblies per contaminated isolate. This procedure resulted in 23 unimodal assemblies, of which two were excluded due to low coverage (), and four were removed because their GC content fell outside or at the edge of the expected *Methylobacterium* range (66.5–72.9%; (Leducq, Sneddon, et al. 2022); Figure S1; Table S2). BLAST analysis of representative scaffold fragments from these four assemblies assigned them to non-*Methylobacterium* taxa (*Stenotrophomonas maltophilia*, *Comamonas* sp., *Hymenobacter* sp.; Table S2), confirming contamination. This step resulted in 79 assemblies.

For six isolates (APS3.4, KUML1, AMHL1, AMHS2, DALS4, GEZL2), we also carried out Oxford nanopore long reads sequencing using the MinION platform (Jain et al. 2016). Hybrid assemblies combining short- and long-read data were generated using Unicycler v0.5.0 in bold mode to optimize assembly completeness (Wick et al. 2017) and showed higher quality compared to short-read-only assemblies, with N50 values ranging from 1,039 to 6,149 kb, compared to 9 to 176 kb for short-read assemblies (Table S2).

The final dataset comprised 79 assemblies, including 73 short-read assemblies and 6 hybrid assemblies. Short-read and long-read sequencing libraries were deposited on NCBI under BioProject PRJNA1546143 (BioSamples SAMN64252281-SAMN64252349). To accommodate multiple assemblies obtained from some libraries, the 79 genome assemblies were deposited on NCBI under a different BioProject (PRJNA1548628; BioSamples SAMN64379562-SAMN64379640; Table S2).

### Methylobacterium genome collection and annotation

We compiled a collection of 293 *Methylobacterium* genomes for comparative analyses (Table S1), comprising 189 previously annotated genomes (Leducq, Sneddon, et al. 2022) plus 104 new *Methylobacterium* assemblies: 79 rice-associated genome assemblies from the present study, 7 complete soybean leaf-associated genomes assemblies (Govindaraju et al. 2025), and 18 recently published genomes including 8 newly described species as of September 2025 (*M. flocculans*, *M. oryzisoli*, *M. alsaeris*, *M. carpenticola*, *M. litchii*, *M. guangdongense*, *M. nigriterrae*, *M. subtropicum* (Chen et al. 2024; Gao et al. 2024; Gao et al. 2025; Liu et al. 2025; Yoon et al. 2025)). All 104 new *Methylobacterium* genomes were annotated individually with RAST (Aziz et al. 2008) as done for the previously annotated genomes (Leducq, Sneddon, et al. 2022).

### Core genome definition

We identified the *Methylobacterium* core genome for phylogenomic inferences and HGT detection as previously (Leducq, Sneddon, et al. 2022) with the following modifications. From the 293 *Methylobacterium* genomes (Table S1), we first removed RAST genes annotated as “hypothetical protein”, “repeat region” and “mobile element protein” and then tallied unique gene annotations.

Because many genome assemblies were draft (incomplete and/or fragmented), we expected a higher number of false duplications/losses for many core genes. In order to maximise the number of genes for which evolutionary history could be compared across diverging lineages, while taking into account lower sequence quality in draft assemblies, we initially set a very relaxed threshold by defining core genes as genes present in at least 80% of assemblies, with an average occurrence of maximum 1.5 copies per genome. Because *Methylobacterium* consists of 4 distinct phylogroups (A, B, C, D (Leducq, Sneddon, et al. 2022)), we applied core genome thresholds separately for each phylogroup. We first defined the core genome from the previous 189 *Methylobacterium* assemblies that had been assigned phylogroup identity (Leducq, Sneddon, et al. 2022), and then included the 104 new assemblies (for which the phylogroup was unknown). We then distinguished true gene duplications from artefactual duplications caused by fragmentation in draft assemblies (single-copy genes split across scaffolds) as follows. First, for each candidate core gene, we estimated median gene size in ‘good’ assemblies (complete or nearly complete genomes, i.e. with N50>0.5Mb); we used this value as the expected size for each gene as previously (Leducq, Sneddon, et al. 2022). Second, for each candidate core gene, we retrieved the gene size of all copies identified across all assemblies and normalized them by the median size in good assemblies as previously (Leducq, Sneddon, et al. 2022): the total of relative sizes for all copies of a given gene within a given assembly should then give a proxy of the actual copy number. Third, we interpreted estimated copy numbers lower than 0.7 as gene deletions, or fragmented copies in draft assemblies, and values above 1.3 as potential gene duplication in some assemblies. We thus considered estimated copy numbers outside this range as missing data, as previously (Leducq, Sneddon, et al. 2022). Finally, to mitigate the effect of gene loss or duplication associated with phylogroups, we removed any candidate core gene with missing data in more than 30% of assemblies within at least one phylogroup. To mitigate biases due to different models of evolution for RNA genes, only protein-encoding genes (PEG) were considered for further analyses.

PEG sequence alignments were performed for each core PEG individually with the R package *msa* (Bodenhofer et al. 2015). Sequence alignments were performed on amino-acid sequences and then back-translated to nucleotide sequences as previously (Leducq, Sneddon, et al. 2022).

### Species and lineage delimitation using ANI

Average nucleotide identity (ANI) was calculated pairwise among the 293 *Methylobacterium* genome assemblies using FastANI (Jain et al. 2018). Genomes were assigned to the same species when their pairwise ANI was ≥95% (Jain et al. 2018). For genomes lacking a type strain reference, new species were defined when their ANI exceeded 95% with at least one other genome in the dataset (Jain et al. 2018). By default, we re-used candidate species’ provisional labeling from our previous work (*Methylobacterium sp. 001* to *044*) and extended it for new candidate species (Leducq, Sneddon, et al. 2022). Species classification was then used to infer the consensus species tree (see section *Phylogenomic inference*).

Within phylogroups A, B, C, and D, lineages were defined to capture finer-scale evolutionary structure relevant for recombination and HGT analyses. Lineages were defined as monophyletic clades in the consensus lineage tree (see *Phylogenomic inference*) and delineation was based on ANI values below the species threshold (<95%) but above phylogroup level (>84%), according to our previous analyses (Alleman et al. 2025).

### Phylogenomic inference

We inferred the consensus *Methylobacterium* lineage and species tree based on core PEGs and accounting for putative HGT and ILS as previously (Leducq, Sneddon, et al. 2022). First, a maximum-likelihood tree was inferred for each PEG individually with RAxML (GTRGAMMA model, 1,000 bootstraps) (Stamatakis 2014). For each core PEG, we visualized the phylogenetic tree using the *plot.phylo()* function from the R package ape (Paradis and Schliep 2019). For each PEG tree, we also retrieved branch length statistics (mean, median, standard deviation and max value) and reported whether each phylogroup (A, B, C, D) and lineage was monophyletic. The consensus lineage tree (each genome treated as a distinct evolutionary unit) and the species tree (genomes from the same species pooled together in the same evolutionary unit) were then inferred by combining the per-PEG trees (nodes with support <10% were collapsed) using quartet-based inferences under the multispecies coalescent model (MSC) accounting for ILS with ASTRAL-IV implemented in ASTER (Zhang et al. 2025).

### KEGG pathway annotation

We used the *M. oryzae* CBMB20 strain genome as a reference, because it corresponded to the phyllosphere *Methylobacterium* strain with the highest genome completeness in our previous study (N50 = 6.29 Mb) (Leducq, Sneddon, et al. 2022). KEGG pathway of protein sequences for each core PEG from the CBMB20 genome was predicted with KASS (Moriya et al. 2007) with default parameters. For the genome database, we used default bacterial genomes and added any genome of any *Methylobacterium* species (including *Methylorubrum* (Alleman et al. 2025)) or closely related Hyphomicrobiales genera (*Beijerinckia*, *Methylocella* (Leducq et al. 2026)) available in the KASS reference database. KEGG pathways classified into “Human Diseases” and “Organismal Systems” were excluded.

### PEG cluster conservation

We estimated average cluster conservation (ACC), i.e. the proportion of pairs of core PEGs consistently colocalizing across diverging *Methylobacterium* genomes. This approach is equivalent to core genome synteny that we previously evaluated in *Methylobacterium* (Leducq, Sneddon, et al. 2022), but less constrained by small-scale genomic rearrangements and by genome incompleteness or fragmentation. For each genome, we reported any possible pair of core PEGs located within 25 kb (conservative threshold; N/A returned if genes were located on different scaffolds or if at least one gene was missing in that genome). ACC among two genomes was calculated as the number of pairs of colocalizing core PEGs they shared relative to the minimum observable number in one genome.

For each possible pair of core PEGs, we also calculated colocalization conservation across *Methylobacterium* lineages (CCL), as the number of lineages where the two PEGs are located within 25 kb, weighted by the number of genomes per species and by the number of species per lineage. This statistic was thereafter used to identify conserved gene clusters (section *Gene clusters and functions significantly enriched in HGT*).

### HGT likelihood using corrected RF distances

To identify core PEGs potentially undergoing HGT during *Methylobacterium* evolution, we hypothesized that HGT of a given gene would result in topological discordance between its phylogenetic tree and the consensus *Methylobacterium* tree. We thus measured the topological divergence between each core PEG phylogenetic tree and the consensus lineage tree using the Robinson-Foulds metric (RF distance) (Robinson and Foulds 1981), implemented using the *RF.dist* function from the R package *phangorn* (Schliep 2011). Phylogeny inference, and thus topology discordance metrics like the RF distance, could also be sensitive to the amount of information available in sequence alignments (number of polymorphic sites) (Philippe et al. 2011), to selection pressures differentially acting on gene evolution (He et al. 2025), and to the number of tips in each gene tree (some low-quality assemblies missing in some gene trees). We thus built a global model to correct RF distances (log-transformed to meet normality) by the number of polymorphic sites per PEG alignment (*nPOL*); by the dN/dS (*dNdS*) ratio per PEG alignment as a proxy for selection, using the R function *kaks* from package *seqinr* (Charif and Lobry 2007); by the number of genome assemblies per PEG tree (*nTIP*); by the median nodal support per PEG tree (*Mns*); and by the interactions between those four factors. We used the following starting model: *lm(log10(RF)* ∼ *nPOL*dNdS*Mns*nTIP)* and iteratively removed non-significant factors and interactions (anova; *p value*>0.05). We extracted residual RF distance values (*RES*) from the best model as a predictor for HGT likelihood.

### KEGG pathway enrichment

We determined KEGG pathways putatively enriched in HGT. For each KEGG pathway, we calculated the average RES value (HGT likelihood) among genes classified within this function. Average observed RES values for each KEGG pathway were compared to expected values after K=1.10^6^ permutations of RES values among PEGs, with nested functions associated to each PEG fixed among permutations. Average RES values were recomputed for each KEGG pathway and each replicate, and two-sided p-values were calculated as pval = (K - Z + 1) / (K + 1) (Z = number of permutation replicates with RES ≥ observed) (Phipson and Smyth 2010). P-values were adjusted to control for false discovery rate using the Benjamini-Hochberg method (Benjamini et al. 2001).

### Methylobacterium lineage evolutionary network

We reconstructed the evolutionary network of *Methylobacterium* combining core PEG tree topologies, simplified at the lineage level, hence focusing on putative gene swaps prior lineage radiation. Considering nodes of the ASTER species tree, simplified at the lineage level, as nodes for our network reconstruction, we first predicted 891 possible edges in the network, corresponding to non-redundant pairwise combinations among those nodes and nested clades (thereafter partitions). Predicted partitions corresponded to the 24 lineages (lineage partitions), to the 21 consensus tree internal branches (consensus partitions), and to 846 partitions that could be interpretated as a single branch swap compared to the consensus tree (alternative partitions), hence putative gene exchanges during *Methylobacterium* evolution. Second, we determined partitions observed across core PEG phylogenetic trees. We selected trees for core PEGs for which sequences could be retrieved from at least 23 out of 24 *Methylobacterium* lineages (970 PEG trees). We removed 129 PEG trees for which information was missing for more than one lineage. When one lineage was missing (case for 259 PEG trees), we artificially added this lineage in the gene tree, conservatively assuming it formed a monophyletic group with its closest lineages according to the consensus ASTER tree. For each tree rooted on the first detected monophyletic lineage (by default lineage A1; function *is.monophyletic* in package *ape* (Paradis and Schliep 2019)), we retrieved all partitions simplified at the lineage level (i.e. nodes containing all genomes from the same lineage(s) were collapsed). For instance, partition A12 indicates that lineages A1 and A2 form a monophyletic group (regardless branch swaps within this groups). We removed partitions that did not include all genomes from a given lineage, hence excluding the detection of potential HGT among lineages before lineage splitting. Third, we determine whether and how frequently the predicted partitions (lineage, consensus and alternative) were observed across the 970 PEG trees. Alternative partitions that were observed but not predicted could not be interpreted as a single branch swap and were excluded from further analyses for simplification. Fourth, we determined whether alternative partitions were enriched for branch swaps within phylogroups (four categories corresponding to swaps within A, B, C or D) compared to swaps between phylogroups (six possible categories corresponding to swaps among A, B, C or D). We performed this analysis on 698 predicted alternative partitions that combined subsets of at most two phylogroups, hence restricting the detection to branch swaps occurring after phylogroup radiation. We tested for differences in proportions of observed partitions compared to expected partitions among categories (4 within and 6 among phylogroups; R function *pairwise_prop_test*). For each category, the retrieved the occurrence of each observed partition among core PEG trees and tested for differences among categories (4 within and 6 among phylogroups) using a Tukey test (R function *TukeyHSD*). Fitht, we tested of observed partition enrichment in KEGG pathways. We performed 100,000 permutations among PEGs (nested functions fixed). The number of each KEGG pathway observed for each partition was recomputed for each replicate, and two-sided p-values were calculated as pval = (K - Z + 1) / (K + 1) (Z = number of replicates with nKEGG ≥ observed) (Phipson and Smyth 2010). P-values were adjusted to control for false discovery rate using the Benjamini-Hochberg method (Benjamini et al. 2001). The *Methylobacterium* evolutionary network was visualized in Cytoscape (Shannon et al. 2003).

### Gene clusters and functions significantly enriched in HGT

We identified gene clusters and functions potentially enriched in HGT. Assuming that bacterial genomes are organized in clusters of genes involved in similar pathways and functions (Fang et al. 2008), we hypothesized 1) that core PEGs involved in the same pathway (KEGGs) are functionally constrained to coevolve and thus to display more similar phylogenetic trees than functionally unrelated genes and 2) that core PEGs colocalizing in *Methylobacterium* genomes are both physically and functionally linked, and thus more likely to display similar phylogenetic trees than unlinked PEGs.

First, we estimated gene cluster size in *Methylobacterium* genomes, i.e. the average distance between two genes likely to experience similar evolutionary constraints. For each possible pair of core PEGs, we calculated 1) *pDIST*, the pairwise distance between gene positions in the CBMB20 reference genomes taking into account chromosome circularity, 2) the number of KEGG pathways shared by each pair of genes (N/A if the information was missing for at least one gene) and 3) the pairwise RF distance (*pRF*) between the phylogenetic trees of each pair of genes. We corrected *pRF* for *nPOL*, *dNdS*, *nTIP* and *Mns* using a global model as described above (see section *HGT likelihood using corrected RF distances*) but including values from both genes of a given pair in the model. We estimated gene cluster size as a function of similarity in gene function (number of shared KEGG pathways, expected to increase with *pDIST*) and phylogenetic dissimilarity (*pRF* expected to decrease with *pDIST*) independently. Briefly, for each discrete kb window of *pDist* in the range 1-100 kb, we calculated average *pRF* and the proportion of gene pairs sharing at least one KEGG pathway (*pKEGG*). Then, we determined the breakpoint in the *pRF*∼*log(pDist)* and in the *pKEGG*∼*log(pDist)* linear regressions (Davies test), and 95% estimates of slopes for each segment of the regression, using the R function *segmented* (Muggeo 2017). To evaluate the effect of gene cluster conservation on PEG coevolution while accounting for genomic rearrangements across diverging *Methylobacterium* lineages (Leducq, Sneddon, et al. 2022), we compared core PEGs colocalization conservation across all lineages (CCL) with their phylogenetic pairwise dissimilarity (*pRF*). To control for data biases toward rare/absent pairs of core PEGs, we tested for the CCL∼*pRF* linear regression both on raw data and on average *pRF* values for 50 discrete CCL binned values.

Second, we identified clusters of core PEGs putatively enriched for HGT. For each discrete CCL value in the range 8-24 lineages (24 meaning that the pair is conserved across all *Methylobacterium* lineages), we aggregated pairs of core PEGs to generate clusters. For each level of conservation in the range 8-24, we estimated HGT likelihood for each cluster. For a given level, we calculated the average RES value among genes classified within the same cluster. The average observed RES value for each cluster was then compared to expected values from K=10^6^ permutations of RES values among genes, and two-sided p-values were calculated as described above. Cluster conservation and HGT enrichment were visualized in Cytoscape (Shannon et al. 2003), with nodes representing core PEGs (colored by RES value) and edges weighted by average pair colocalization across lineages (option *prefuse force-directed layout*).

### Detection of horizontal gene transfer using nucleotide polymorphism patterns

We detected HGT based on nucleotide polymorphism patterns, an approach complementary to phylogenetic discordance, using hierarchical Bayesian Analysis of Population Structure (BAPS) implemented in hierBAPS R package (Tonkin-Hill et al. 2018), which identifies genetic structure from multiple sequence alignments without relying on gene tree topologies. Each of the 1,099 core PEGs was analyzed independently, based on the same nucleotide sequence alignments used for phylogenetic inference (see section *Core-genome definition*). BAPS did not consider recombination within genes, which we assumed to be negligible in *Methylobacterium* core PEG sequences compared to recombination among them (Leducq, Sneddon, et al. 2022). For each core PEG, each sequence was assigned to a BAPS *lineage*. We first determined the number of populations (nPOP) from which the number of BAPS lineages would stabilize for a subset of 190 core PEGs, assuming number of populations in the range 2-60. The average number of BAPS lineages inferred from a subset of 190 core PEGs and the average proportion of BAPS lineages per PEG that matched the corresponding gene tree topology started to stabilize when assuming approximately 20-30 ancestral populations, broadly corresponding to the number of *Methylobacterium* lineages (*n* = 24; Figure S10A-B), and this value was subsequently retained for all analyses. Thereafter, BAPS lineage assignments were processed to all core PEGs to assess congruence with the consensus phylogeny and predefined phylogroups (A–D). For each core PEG, BAPS lineages were re-annotated *a posteriori* according to the phylogroup membership of the strains they contained. BAPS lineages composed exclusively of strains from a single phylogroup and forming a monophyletic clade in both the consensus species tree and the corresponding gene tree were interpreted as no evidence of HGT. BAPS lineages composed of strains from a single phylogroup, congruent with the corresponding gene tree, but incongruent with the consensus species tree topology, were classified as intra-phylogroup HGT, consistent with recombination among closely related lineages. In contrast, BAPS lineages containing strains from two or more distinct phylogroups, congruent with the corresponding gene tree, were classified as inter-phylogroup HGT, reflecting putative horizontal transfer across deeper phylogenetic boundaries. In some cases, BAPS lineages could not be reliably reconciled with either of the individual gene tree or the consensus tree, typically due to limited nucleotide polymorphism or low phylogenetic resolution for some PEGs. These cases were conservatively classified as undetermined. For each core PEGs, we reported the proportion of BAPS lineages corresponding to putative HGT (undetermined cases removed) and tested for its correlation with HGT likelihood (RES) with a linear model.

### Bacteriochlorophyll and carotenoid biosynthesis gene cluster evolution and structure analysis in Methylobacterium and closely related Hyphomicrobiales

We evaluated the evolution of gene sequences and cluster organization of the bacteriochlorophyll and carotenoid gene (BCHL) cluster. To detect potential HGT from outside *Methylobacterium*, we extended the BCHL cluster analysis to closely related Hyphomicrobiales genomes, including groups *RH_AL1* and Lichenihabitantaceae inhabiting the phyllosphere (Leducq et al. 2026). First, we re-inferred the phylogeny for the 32 *Methylobacterium* core PEGs associated with the BCHL cluster (BCHL genes; Table 1), including gene copies identified across the 293 *Methylobacterium* genomes, and the 167 annotated genomes and metagenome-assembled genomes (MAGs) of related Hyphomicrobiales from our previous work (Leducq, Sneddon, et al. 2022; Leducq et al. 2026). Nucleotide sequences of the 32 BCHL genes were retrieved from RAST annotation files available from our previous works (Leducq, Sneddon, et al. 2022; Leducq et al. 2026), and alignments and phylogenies were generated and visualized as described in the sections *Core-genome definition* and *Phylogenomic inference*. Because the *Methylobacterium* core genome was defined with relaxed criteria, and because variations in copy number of some *Methylobacterium* core PEGs are expected for other Hyphomicrobiales genomes, all gene copies identified for the 32 BCHL genes were included in phylogenetic analyses, regardless of copy number.

Second, we compared the BCHL cluster architecture (gene organization) across *Methylobacterium* lineages and related Hyphomicrobiales genera with identified BCHL clusters. To limit biases due to fragmented genomes and MAGs included in our dataset, we restricted this analysis to Hyphomicrobiales genomes and MAGs with at least 25 BCHL genes, corresponding to the minimum necessary to conserve at least one genome per genus with a BCHL cluster. For *Enterovirga*, *JAJXWB01* and *Rhabdaerophilum*, only fragmented MAGs were available, and the BCHL architecture could not be assessed. For each selected genome and MAG, BCHL gene positions (scaffold, start, end) were retrieved, and BCHL genes were aggregated in sub-clusters when located in the same scaffold and within a window of 25 kb, as previously (section *Gene clusters and functions significantly enriched in HGT*). For MAGs and highly fragmented genomes, *bchG* and *bchH* positions inside or outside the BCHL cluster were manually checked in the RAST output file.

Third, diversity in BCHL cluster organization was evaluated at both the species and lineage levels for *Methylobacterium* only. For each *Methylobacterium* species and lineage, only the genome with the highest assembly quality (as measured by N50) was retained to avoid redundancy. BCHL cluster architecture was summarized as the number and size of subclusters per genome, ordered according to the consensus species tree (see section *Phylogenomic inference*).

Fourth, BCHL cluster analyses were refined for comparative gene cluster alignment using clinker (Gilchrist and Chooi 2021), enabling visualization and assessment of gene order conservation, rearrangements, and gene content variability across lineages. To limit the number of clusters to display in clinker (because of large memory usage), *Methylobacterium* and other Hyphomicrobiales BCHL clusters were analyzed separately, while including four representative *Methylobacterium* genomes (one from each phylogroup) in the Hyphomicrobiales analysis to allow downstream comparisons. For each clinker analysis, sequences of BCHL subclusters (including intergenic regions) from the best-quality genome of each *Methylobacterium* lineage (n = 24) or all selected BCHL subclusters from Hyphomicrobiales genera (n = 18 + 4 *Methylobacterium*) were pooled into a single FASTA file and annotated *de novo* using the RAST server. Output annotated GenBank (GBK) files were parsed and reorganized so that subclusters from the same genome were grouped into lineage-specific GBK files. These files were then used for clinker analysis, with default parameters.

To display changes in the BCHL cluster structure in an evolutionary context, we inferred a consensus Hyphomicrobiales phylogeny combining this study (*Methylobacterium* lineages) and a phylogenetic tree of closely related Hyphomicrobiales families previously built according to the same pipeline (core gene ML phylogenies individually inferred with RAxML and then combined with ASTER), but on a smaller subset of core genes (n = 824) identified across Hyphomicrobiales genomes and metagenomes (Leducq et al. 2026). Lineage trees from both studies were manually combined in the same newick tree file as follows: in the Hyphomicrobiales tree, we replaced the *Methylobacterium* clade with the lineage tree inferred in the current study. The tree was then rooted on *Rhabdaerophillum* and displayed in FigTree 1.4.4.

## Acknowledgments

The computations were partly conducted using the VALERIA research computing infrastructure at Université Laval (https://valeria.science/). We would like to thank Jerome Laroche, Remi Maglione, Jonathan Jacobs and Hannah Toth for discussions, Shreya Vichare for laboratory assistance, and the NCBS NGS facility for sequencing.

## Data and code availability statements

Code and data are avalaible on github (https://github.com/JBLED/HGT_Methylobacterium_Core_Genome) and Zenodo (https://doi.org/10.5281/zenodo.23768560). Raw sequencing reads and genomes assemblies were deposited in NCBI under BioProjects PRJNA1546143 and PRJNA1548628, respectively.

