## Supplementary material for "Life on leaves left a lasting mark on the *Methylobacterium* core genome": Figure S1

**Figure S1**: Plot of coverage (log scale) against GC content per scaffold for 11 contaminated assemblies from short-read (SR) libraries. A) MANGS3-SR20; B) APS3.4-SR18; C) CONTS2; D) MANL2.2; E) PMS2; F) PNL1; G) GEGL2; H) TPS; I) AMKS5; J) PMPHAS6; K) TAOS4. Each scaffold is represented by a dot (size proportional to scaffold size). Scaffolds shorter than 1 kb were filtered out (in red). Scaffold larger than 1 kb (green) were spliced in different assemblies according to threshold on coverage (dotted grey line). Resulting assemblies were labelled a, b or c. Assemblies with average coverage lower than 6x were not considered in further analyses (red). Four assemblies were excluded because different from *Methylobacterium* (purple).

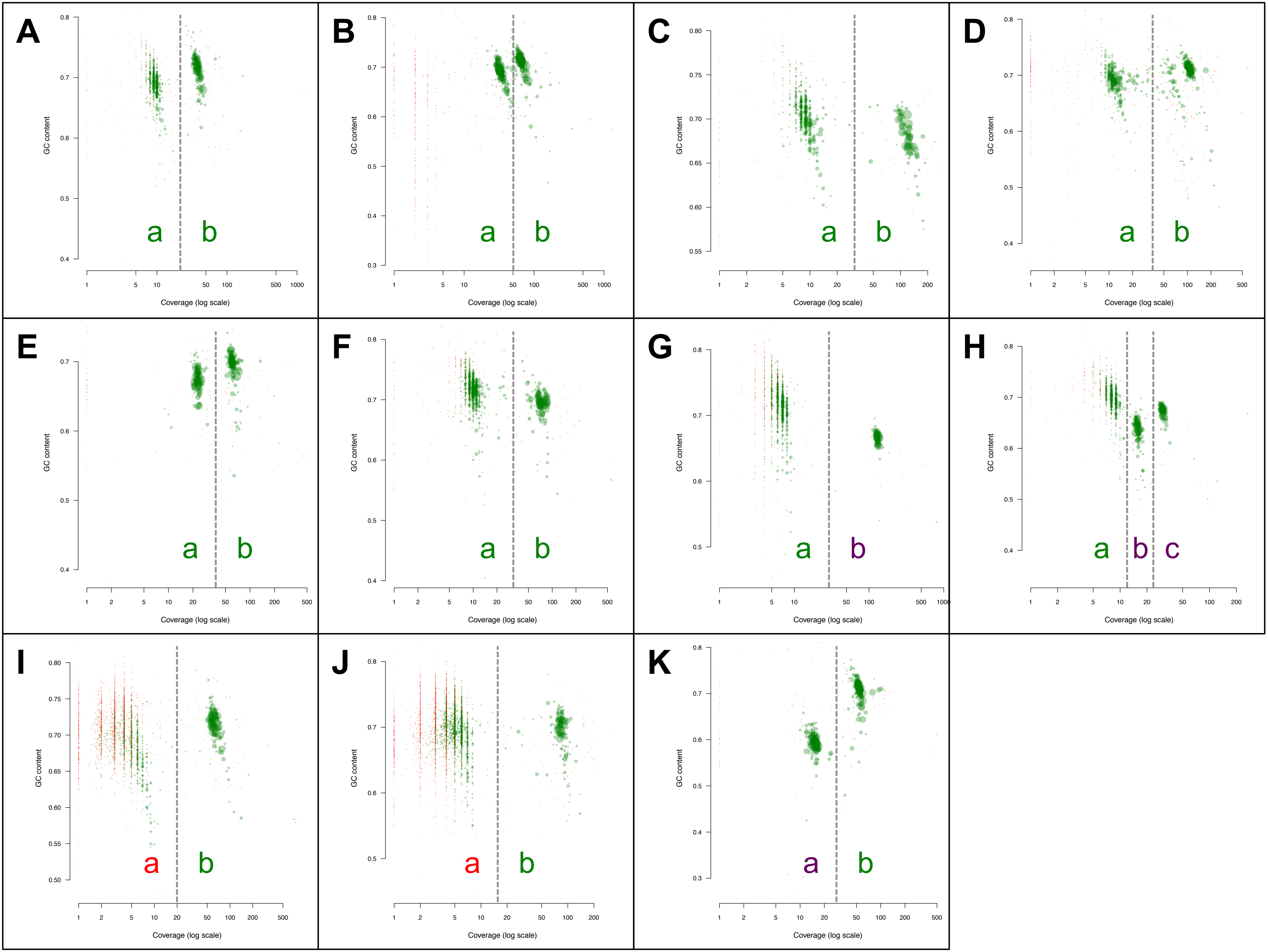

A
GC content
Coverage (log scale)
a
b
B
GC content
Coverage (log scale)
a
b
C
GC content
Coverage (log scale)
a
b
D
GC content
Coverage (log scale)
a
b
E
GC content
Coverage (log scale)
a
b
F
GC content
Coverage (log scale)
a
b
G
GC content
Coverage (log scale)
a
b
H
GC content
Coverage (log scale)
a
b
c
I
GC content
Coverage (log scale)
a
b
J
GC content
Coverage (log scale)
a
b
K
GC content
Coverage (log scale)
a
b
