## Supplementary material for "Life on leaves left a lasting mark on the *Methylobacterium* core genome": Figure S2

**Figure S2:** Number of core genes retrieved per genome assembly, in function of N50 (kb, log scale). Colors indicate genomes assignment to *Methylobacterium* phylogroups from previously (A: red, B: purple; D: blue; C: green(Leducq et al. 2026)) or new assemblies in this study (yellow)

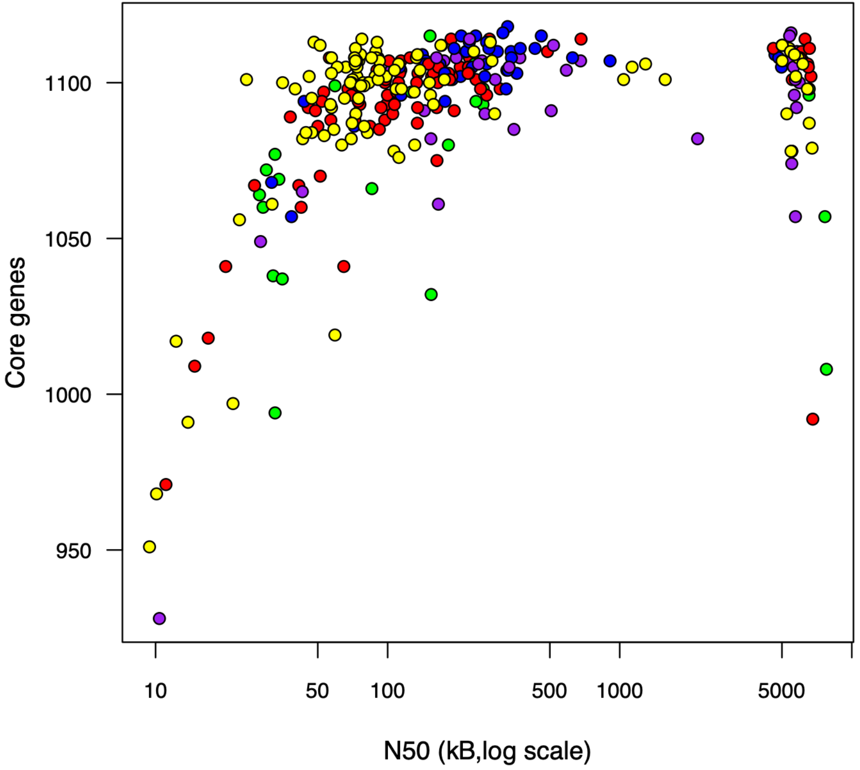

Core genes
1100
1050
1000
950
10
50
100
500
1000
5000
N50 (kB,log scale)
