## Supplementary material for "Life on leaves left a lasting mark on the *Methylobacterium* core genome": Figure S3

**Figure S3:** Distribution of ANI among *Methylobacterium* genomes. Red dotted lines indicate cut-off thresholds that were tested for lineage delineation.

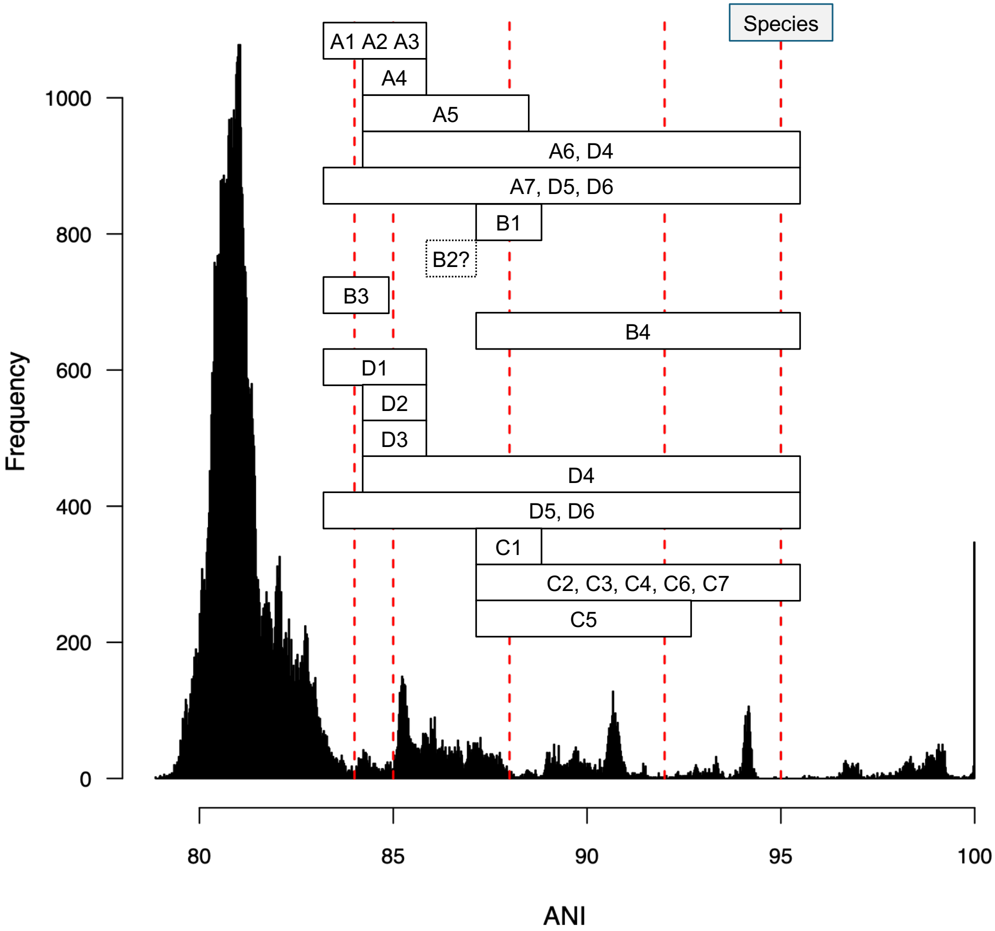

Species
A1 A2 A3
A4
A5
A6, D4
A7, D5, D6
B1
B2?
B3
B4
D1
D2
D3
D4
D5, D6
C1
C2, C3, C4, C6, C7
C5
Frequency
1000
800
600
400
200
0
80
85
90
95
100
ANI
