## Supplementary figures and images for "Life on leaves left a lasting mark on the *Methylobacterium* core genome"

### Figure S4

**Figure S4:** Detailed ASTER lineage tree

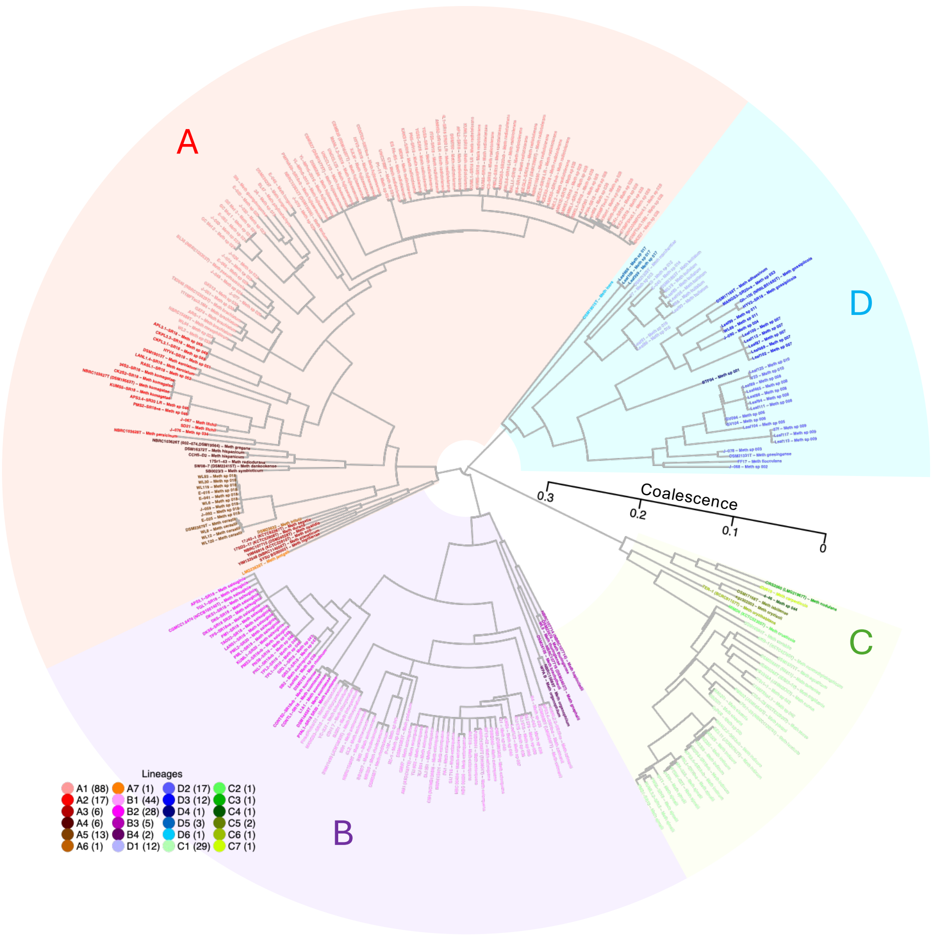
A
D
C
B
Coalescence
0.3
0.2
0.1
0
Lineages
A1 (88)
A2 (17)
A3 (6)
A4 (6)
A5 (13)
A6 (1)
A7 (1)
B1 (44)
B2 (28)
B3 (5)
B4 (2)
D1 (12)
D2 (17)
D3 (12)
D4 (1)
D5 (3)
D6 (1)
C1 (29)
C2 (1)
C3 (1)
C4 (1)
C5 (2)
C6 (1)
C7 (1)
