## Supplementary material for "Life on leaves left a lasting mark on the *Methylobacterium* core genome": Figure S5

**Figure S5:** Estimation of gene cluster size in *Methylobacterium* genome assuming coevolution between pairs of colocalizing genes. A) Average pairwise phylogenetic dissimilarity between genes (pRF) and B) proportion of genes pairs sharing KEGG pathways (pKEGG) were calculated per discrete windows of 1 kb along the CBMB20 genome (distance separating genes; x-axis, log scale). C) Core gene colocalizing in *Methylobacterium* genomes (x-axis) have in average lower phylogenetic dissimilarity (y-axis). For each possible pair of protein-encoding core genes (1,099 genes; 113,169 pairs), Conservation of gene colocalization across lineages (CCL: number of lineages with genes found within the same 25 kb region; x-axis) in function of their pairwise phylogenetic dissimilarity (pRF; y-axis). For graphical purpose, and because pairwise comparisons were biased toward unrelated genes, data were averaged by bins (50 bins; 200-200,000 pairwise comparisons per bin). Linear regressions were significant for both raw data (red line; STAT TEST NEEDED) and binned data (blue line; STAT TEST NEEDED). Open circles indicate data averaged by bins for graphical purposes (50 bins; 200-200,000 pairwise comparisons per bin; linear regression: blue line).

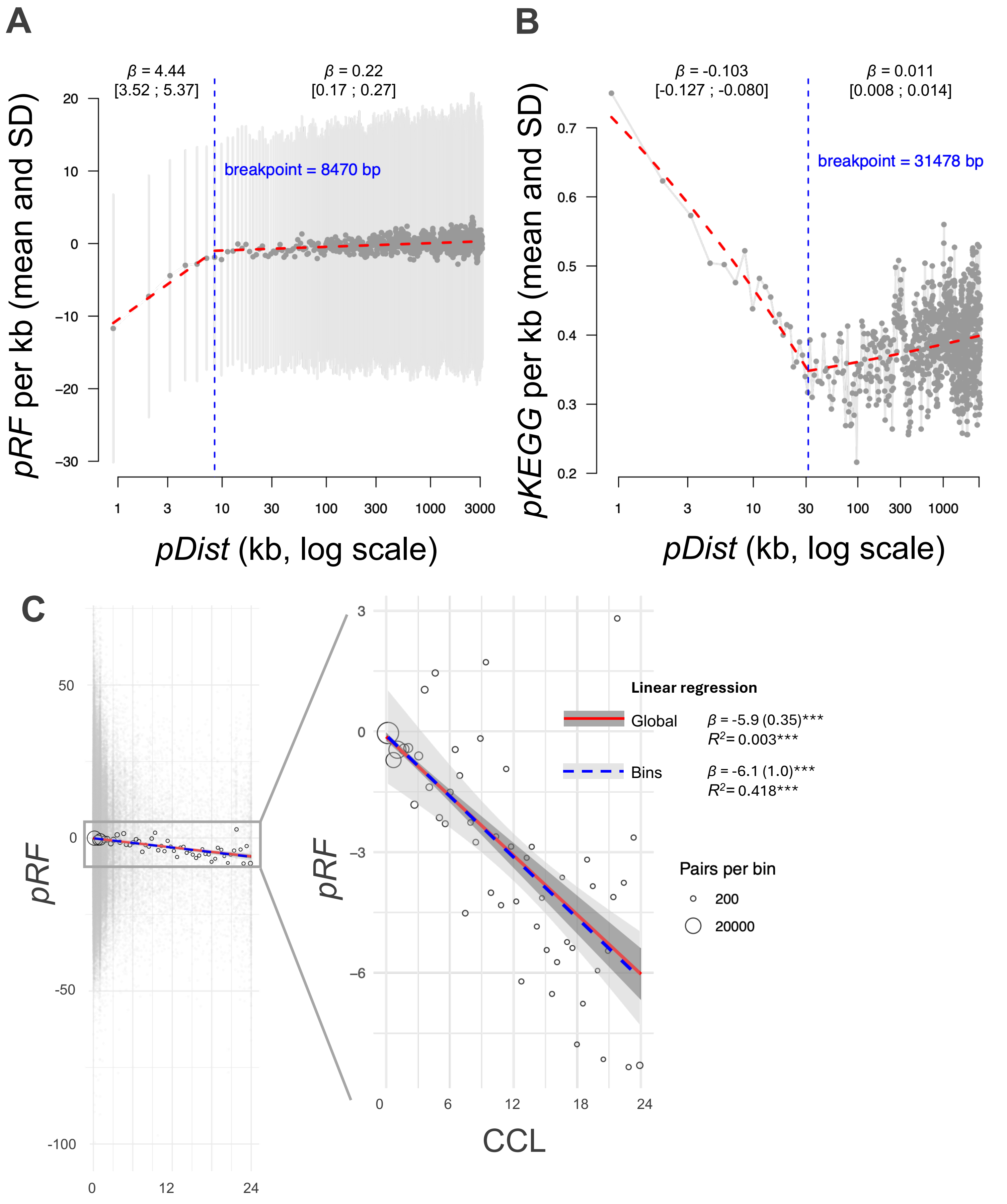

A
β = 4.44
[3.52 ; 5.37]
β = 0.22
[0.17 ; 0.27]
breakpoint = 8470 bp
pRF per kb (mean and SD)
pDist (kb, log scale)
B
β = -0.103
[-0.127 ; -0.080]
β = 0.011
[0.008 ; 0.014]
breakpoint = 31478 bp
pKEGG per kb (mean and SD)
pDist (kb, log scale)
C
pRF
Linear regression
Global
β = -5.9 (0.35)***
R²= 0.003***
Bins
β = -6.1 (1.0)***
R²= 0.418***
Pairs per bin
200
20000
pRF
CCL
