## Supplementary material for "Life on leaves left a lasting mark on the *Methylobacterium* core genome": Figure S6

**Figure S6:** Pairwise ANI and ACC values among *Methylobacterium* genomes are significantly correlated (F-test; quadratic regression). Each point represents a pairwise comparison between two genome assemblies, shape and color according to whether genomes are from the same species (black) or not (grey). Average pairwise comparisons among genomes from the same lineage are indicated.

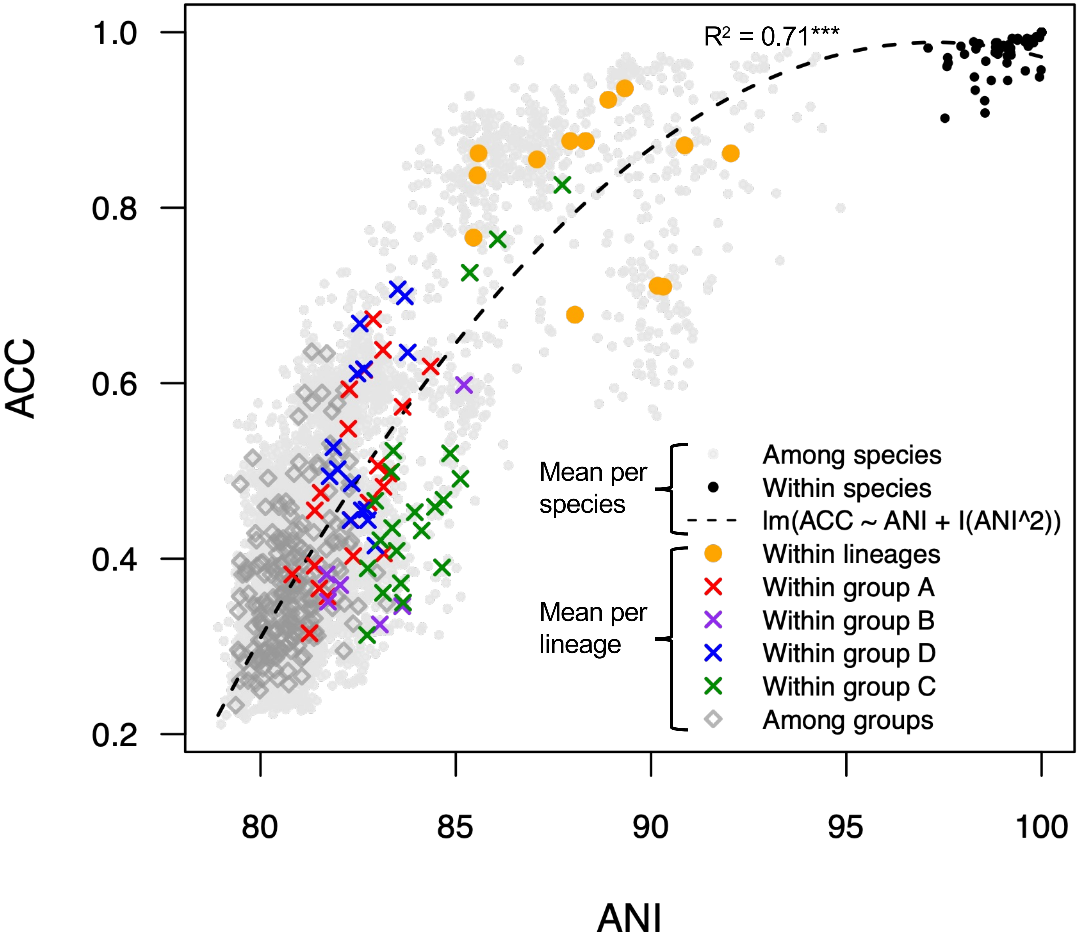

R² = 0.71***
ACC
ANI
1.0
0.8
0.6
0.4
0.2
80
85
90
95
100
Mean per species
Among species
Within species
lm(ACC ~ ANI + I(ANI^2))
Mean per lineage
Within lineages
Within group A
Within group B
Within group D
Within group C
Among groups
