## Supplementary material for "Life on leaves left a lasting mark on the *Methylobacterium* core genome": Figure S7

**Figure S7:** Correction of RF distances between 1,099 PEG trees and the consensus lineage tree. A) RF in function of *dNdS*; B) of *nPOL;* C) of *Mns* and D) of *TIP*. Left panel show regression with uncorrected RF distance. Right panels show regressions with corrected RF distance, corresponding to residuals values (RES) of the best linear regression model (Table S7).

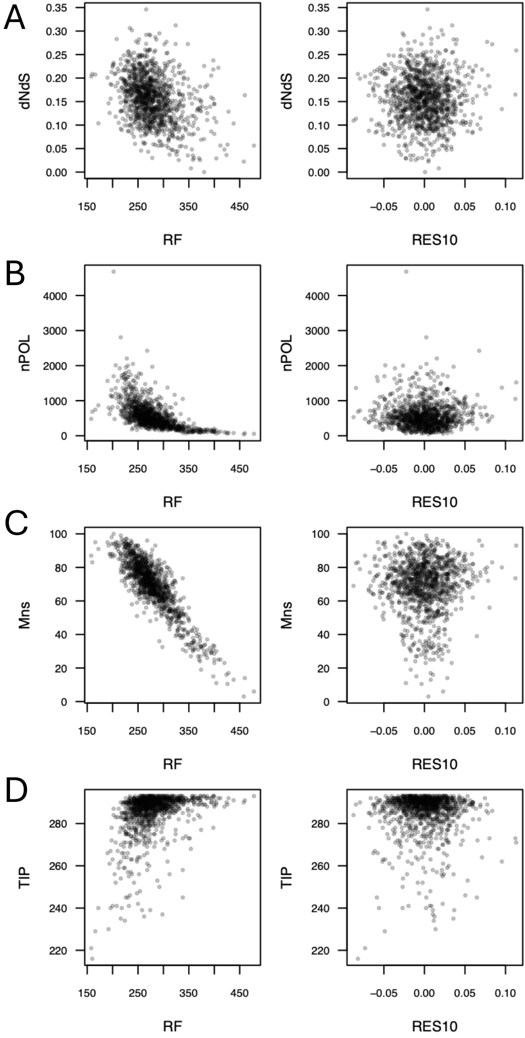
A
B
C
D
dNdS
nPOL
Mns
TIP
RF
RES10
0.00
0.05
0.10
0.15
0.20
0.25
0.30
0.35
0
1000
2000
3000
4000
20
40
60
80
100
220
240
260
280
150
250
350
450
−0.05
0.00
0.05
0.10
