## Supplementary material for "Life on leaves left a lasting mark on the *Methylobacterium* core genome": Figure S8

**Figure S8:** Average residual RF distance per KEGG pathway showing that genes involved in porphyrin metabolism (bacteriochlorophyll precursor) and carotenoid biosynthesis have significantly higher residual RF (y-axis) value than expected by chance (x-axis; red; $p < 0.001$; 100,000 permutation, BH correction).

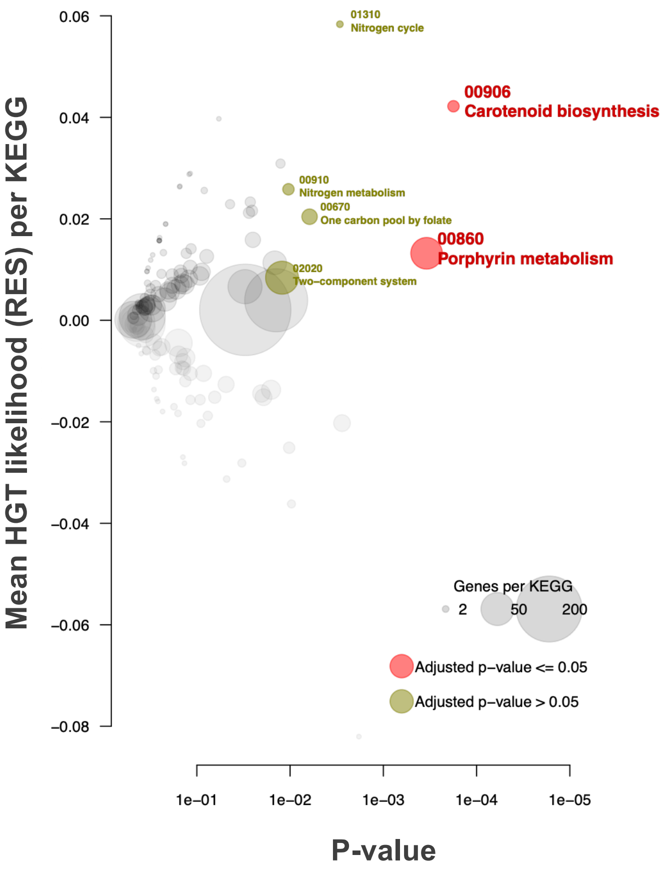

Mean HGT likelihood (RES) per KEGG
0.06
0.04
0.02
0.00
−0.02
−0.04
−0.06
−0.08
01310
Nitrogen cycle
00906
Carotenoid biosynthesis
00910
Nitrogen metabolism
00670
One carbon pool by folate
00860
Porphyrin metabolism
02020
Two−component system
Genes per KEGG
2
50
200
Adjusted p−value <= 0.05
Adjusted p−value > 0.05
1e−01
1e−02
1e−03
1e−04
1e−05
P-value
