## Supplementary material for "Life on leaves left a lasting mark on the *Methylobacterium* core genome": Figure S9

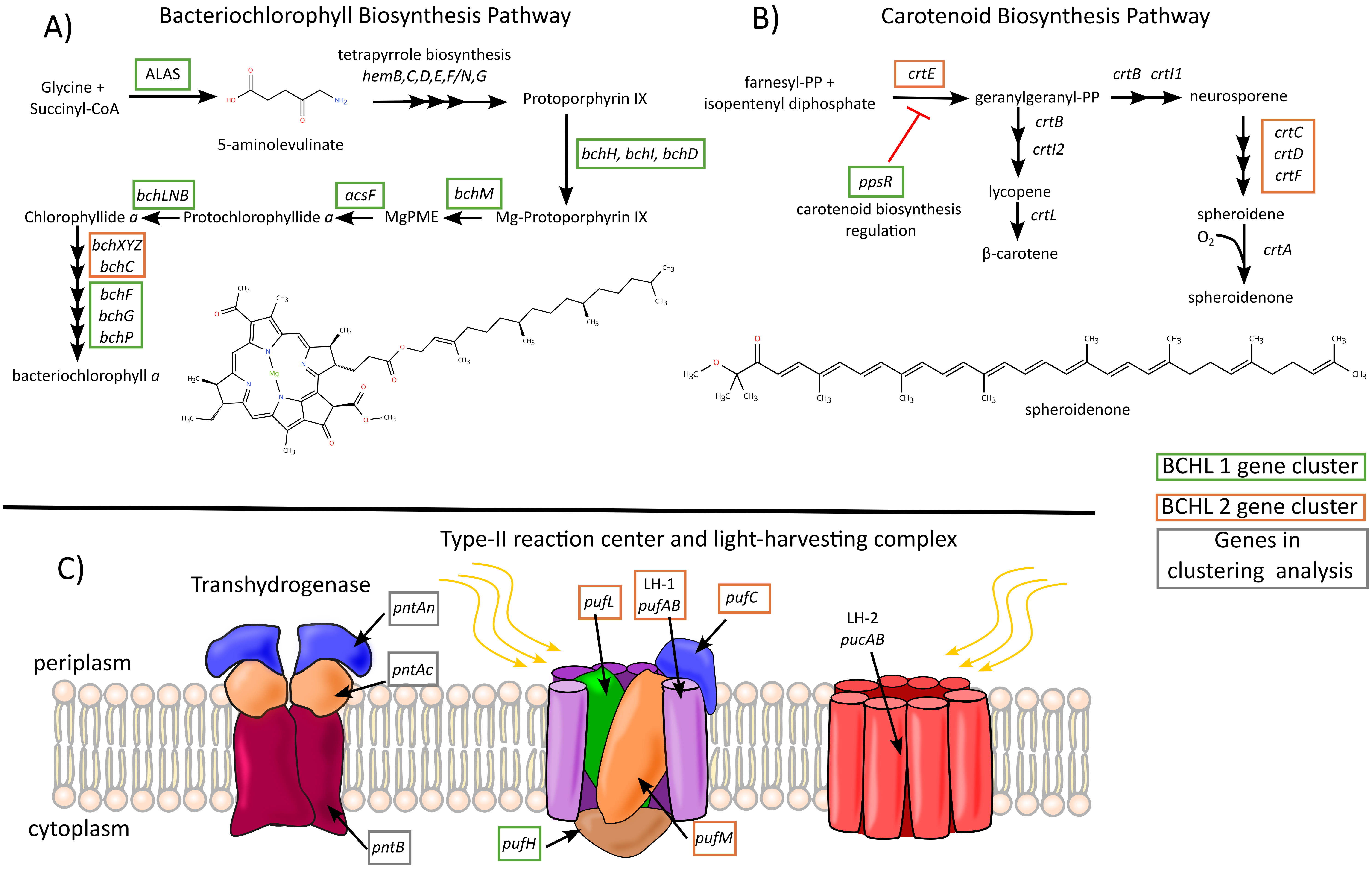
A)
Bacteriochlorophyll Biosynthesis Pathway
Glycine +
Succinyl-CoA
ALAS
5-aminolevulinate
tetrapyrrole biosynthesis
hemB,C,D,E,F/N,G
Protoporphyrin IX
bchH, bchI, bchD
Mg-Protoporphyrin IX
bchM
MgPME
acsF
Protochlorophyllide a
bchLNB
Chlorophyllide a
bchXYZ
bchC
bchF
bchG
bchP
bacteriochlorophyll a
B)
Carotenoid Biosynthesis Pathway
farnesyl-PP +
isopentenyl diphosphate
crtE
geranylgeranyl-PP
crtB crtI1
neurosporene
ppsR
carotenoid biosynthesis
regulation
crtB
crtI2
lycopene
crtL
β-carotene
crtC
crtD
crtF
spheroidene
O2
crtA
spheroidenone
spheroidenone
BCHL 1 gene cluster
BCHL 2 gene cluster
Genes in
clustering analysis
C)
Type-II reaction center and light-harvesting complex
Transhydrogenase
periplasm
cytoplasm
pntAn
pntAc
pntB
pufL
LH-1
pufAB
pufC
pufH
pufM
LH-2
pucAB

**Figure S10:** Summary of BAPS analyses across *Methylobacterium* core PEGs. (A) Number of BAPS lineages and (B) proportion of BAPS lineages that are conserved for analysis after removing incongruences with the gene tree, both estimated in the range 2-60 populations assumed for the BAPS analysis of 190 core PEG trees. Blue lines indicate individual PEG analyses. The black line indicates mean value over 190 PEGs. C) For each *Methylobacterium* lineage (n=24) number of PEGs supporting HGT within (red) or among (yellow) phylogroups. Average values calculated among species from a same lineage. D) Venn diagram showing the number of PEGs with at least one evidence of HGT within (red) and/or among phylogroups (yellow). Numbers are indicated for either all PEGs (n=1,099), and gene clusters enriched in putative HGT (KDP, BCHL).
