## Supplementary material for "Life on leaves left a lasting mark on the *Methylobacterium* core genome": Figure S10

**Figure S10:** Summary of BAPS analyses across *Methylobacterium* core PEGs. (A) Number of BAPS lineages and (B) proportion of BAPS lineages that are conserved for analysis after removing incongruences with the gene tree, both estimated in the range 2-60 populations assumed for the BAPS analysis of 190 core PEG trees. Blue lines indicate individual PEG analyses. The black line indicates mean value over 190 PEGs. C) For each *Methylobacterium* lineage (n=24) number of PEGs supporting HGT within (red) or among (yellow) phylogroups. Average values calculated among species from a same lineage. D) Venn diagram showing the number of PEGs with at least one evidence of HGT within (red) and/or among phylogroups (yellow). Numbers are indicated for either all PEGs (n=1,099), and gene clusters enriched in putative HGT (KDP, BCHL).

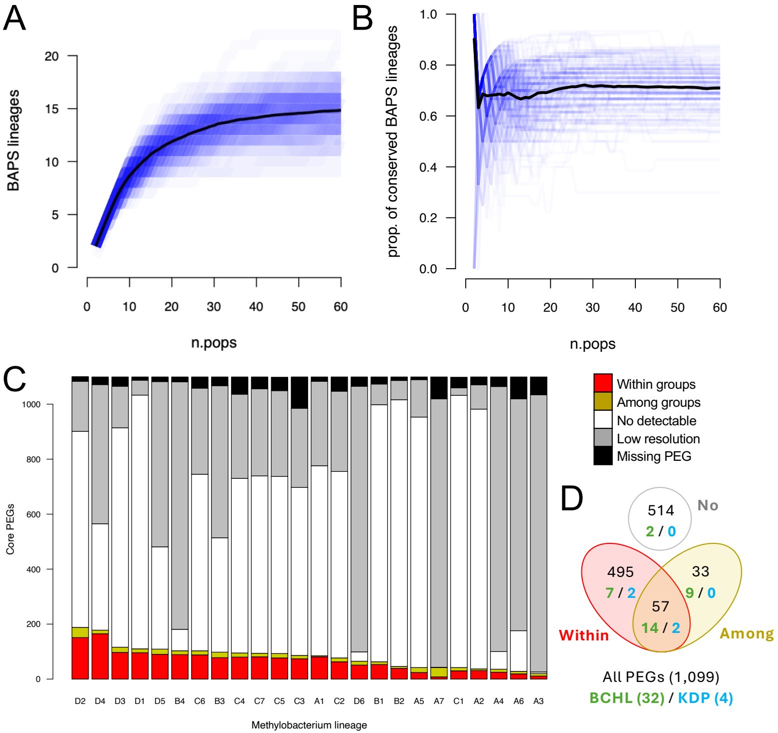

A
BAPS lineages
n.pops
B
prop. of conserved BAPS lineages
n.pops
C
Core PEGs
Methylobacterium lineage
D2 D4 D3 D1 D5 B4 C6 B3 C4 C7 C5 C3 A1 C2 D6 B1 B2 A5 A7 C1 A2 A4 A6 A3
Within groups
Among groups
No detectable
Low resolution
Missing PEG
D
514
2 / 0
No
495
7 / 2
33
9 / 0
57
14 / 2
Within
Among
All PEGs (1,099)
BCHL (32) / KDP (4)
