## Supplementary material for "Life on leaves left a lasting mark on the *Methylobacterium* core genome": Figure S11

**Figure S11.** BCHL cluster architecture in *Methylobacterium* species (A) and lineages (B). For each species or lineage (in row, aligned according to the species tree), results are displayed for the genome with the best assembly (higher N50). For each BCHL gene (in column), colors indicate whether gene is absent (black), located in one of the two subclusters (orange or green; shade proportional to subcluster fragmentation), located in a large cluster (purple) or outside clusters (red).

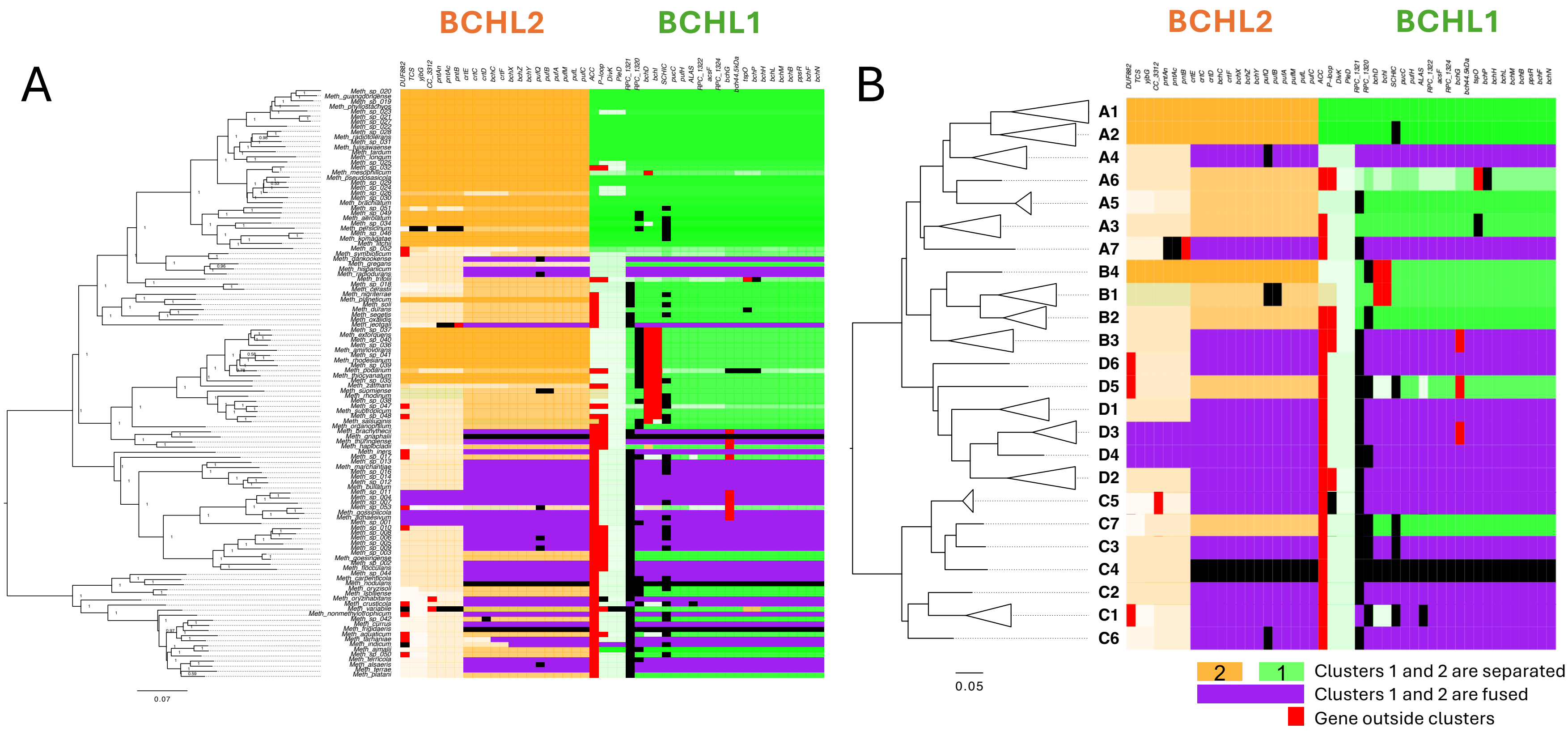

A
B
BCHL2
BCHL1
BCHL2
BCHL1
0.07
0.05
A1
A2
A4
A6
A5
A3
A7
B4
B1
B2
B3
D6
D5
D1
D3
D4
D2
C5
C7
C3
C4
C2
C1
C6
2
1
Clusters 1 and 2 are separated
Clusters 1 and 2 are fused
Gene outside clusters
