## Supplementary material for "Life on leaves left a lasting mark on the *Methylobacterium* core genome": Figure S12

**Figure S12:** Detailed Clinker BCHL cluster architecture in *Methylobacterium* lineages. For each lineage (in row, aligned according to the species tree, collapsed by lineage), results are displayed for the genome with the best assembly (higher N50). Genes are represented by arrows (size proportional to gene length), aligned according to protein sequence similarity among lineages (grey frames when gene sequence homology above 30%). Gene position in BCHL subclusters (green and orange) are schematized in the top panel (see legend for number correspondence with gene names).

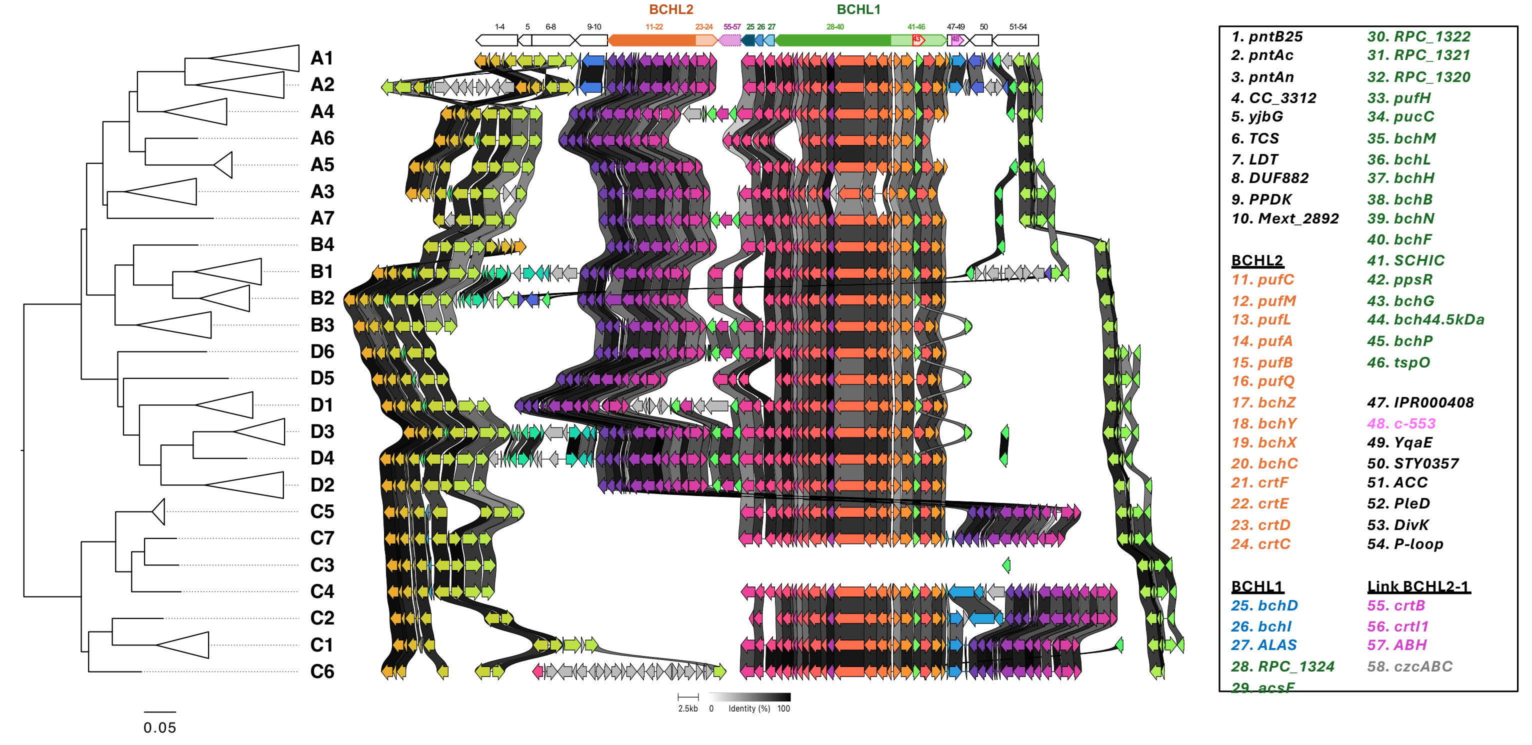

BCHL2
BCHL1
1-4
5
6-8
9-10
11-22
23-24
55-57
25
26
27
28-40
41-46
47-49
50
51-54
43
48
A1
A2
A4
A6
A5
A3
A7
B4
B1
B2
B3
D6
D5
D1
D3
D4
D2
C5
C7
C3
C4
C2
C1
C6
0.05
2.5kb
0
Identity (%)
100
1. pntB25
2. pntAc
3. pntAn
4. CC_3312
5. yjbG
6. TCS
7. LDT
8. DUF882
9. PPDK
10. Mext_2892
BCHL2
11. pufC
12. pufM
13. pufL
14. pufA
15. pufB
16. pufQ
17. bchZ
18. bchY
19. bchX
20. bchC
21. crtF
22. crtE
23. crtD
24. crtC
BCHL1
25. bchD
26. bchI
27. ALAS
28. RPC_1324
29. acsF
30. RPC_1322
31. RPC_1321
32. RPC_1320
33. pufH
34. pucC
35. bchM
36. bchL
37. bchH
38. bchB
39. bchN
40. bchF
41. SCHIC
42. ppsR
43. bchG
44. bch44.5kDa
45. bchP
46. tspO
47. IPR000408
48. c-553
49. YqaE
50. STY0357
51. ACC
52. PleD
53. DivK
54. P-loop
Link BCHL2-1
55. crtB
56. crtI1
57. ABH
58. czcABC
