## Supplementary material for "Life on leaves left a lasting mark on the *Methylobacterium* core genome": Figure S13

**Figure S13:** Detailed Clinker BCHL cluster architecture in four representative *Methylobacterium* genomes (one per phylogroup) and 18 genomes from closely related Hyphomicrobiales genera with complete BCHL cluster (*Salinarimonas*, *Bosea*, *Roseiarcus*, *Rhodoblastus*, *Methylocystis*, *Methylocella*, *Methylocapsa*, *RH_AL1*, *Lichenifustis*, *Lichenihabitans* and *Lichenibacterium*). Genes are represented by arrows (size proportional to gene length), aligned according to protein sequence similarity among genomes (grey frames when gene sequence homology above 30%). Gene position in BCHL subclusters (green and orange) are schematized in the top panel (see legend for number correspondence with gene names in figure S11).

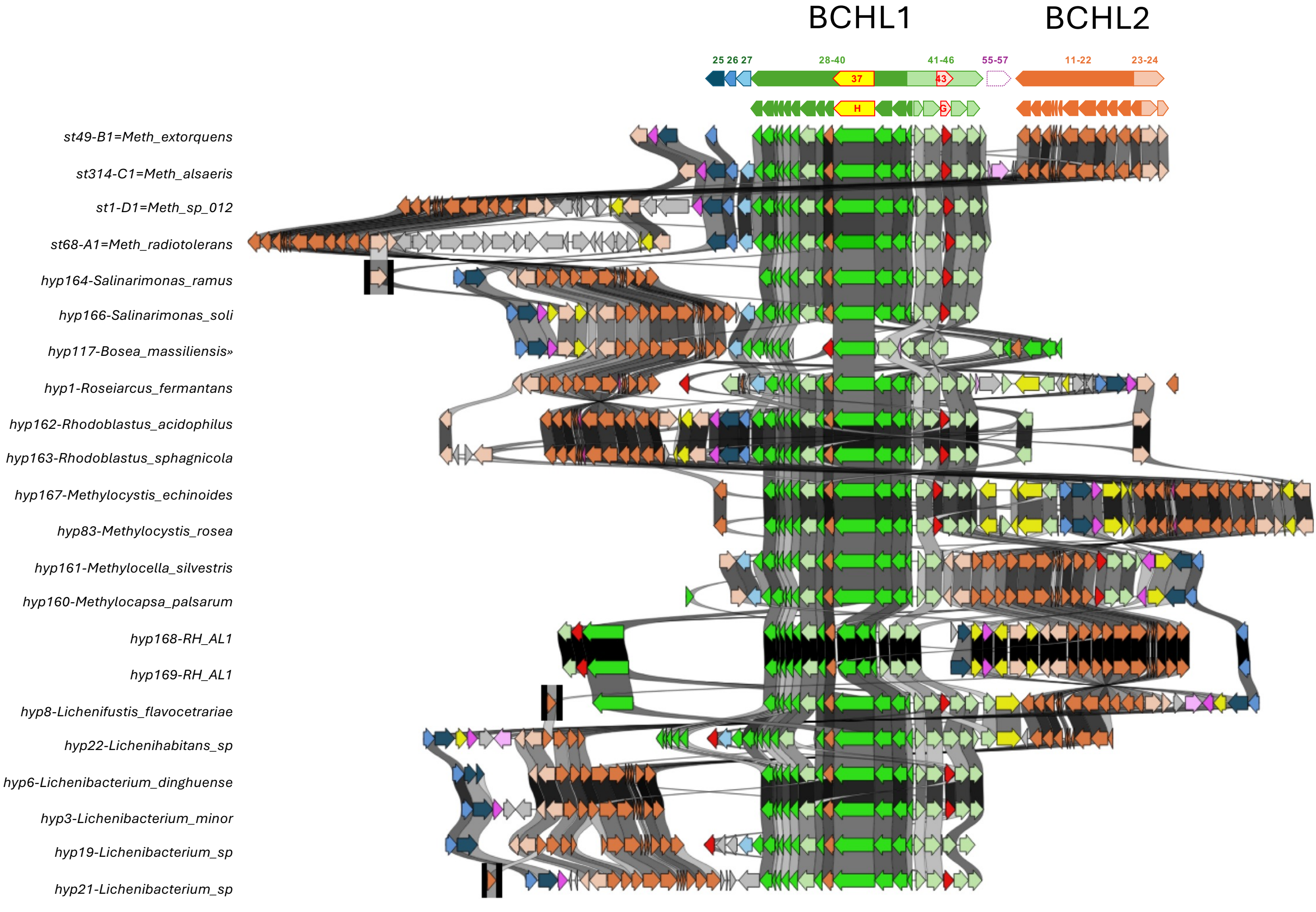

BCHL1
BCHL2
25 26 27
28-40
41-46
55-57
11-22
23-24
37
43
H
G
st49-B1=Meth_extorquens
st314-C1=Meth_alsaeris
st1-D1=Meth_sp_012
st68-A1=Meth_radiotolerans
hyp164-Salinarimonas_ramus
hyp166-Salinarimonas_soli
hyp117-Bosea_massiliensis»
hyp1-Roseiarcus_fermantans
hyp162-Rhodoblastus_acidophilus
hyp163-Rhodoblastus_sphagnicola
hyp167-Methylocystis_echinoides
hyp83-Methylocystis_rosea
hyp161-Methylocella_silvestris
hyp160-Methylocapsa_palsarum
hyp168-RH_AL1
hyp169-RH_AL1
hyp8-Lichenifustis_flavocetrariae
hyp22-Lichenihabitans_sp
hyp6-Lichenibacterium_dinghuense
hyp3-Lichenibacterium_minor
hyp19-Lichenibacterium_sp
hyp21-Lichenibacterium_sp
