## Supplementary material for "Life on leaves left a lasting mark on the *Methylobacterium* core genome": Figure S14

**Figure S14:** Detailed phylogenies of complete gene copies of *pufC* (A), *pufL* (B) and *pufM* (C) encoding for photosynthetic reaction center. For each gene phylogeny (bottom panel), the core *Methylobacterium pufC1*, *pufL1* and *pufM1* clades are indicated in red; extra clades *pufC2*, *pufL2* and *pufM2* observed in some genomes of lineages A1 and A2, in some *Lichenibacterium* and *RH_AL1* genomes are indicated in blue, extra clade *pufC3* observed in somes genomes of lineage A3 and in *RH_AL1* , and clades *pufL3* and *pufM3* observed in *RH_AL1* are indicated in yellow; extra clades observed elsewhere in Lichenihabitantaceae are indicated in grey (*pufC4*, *pufL4*, *pufL4*/5*).* For each gene, relative abundance of each clade combination observed in *Methylobacterium* lineages A1, A2, A3, RH_AL1 and Lichenihabitantaceae genera are indicated in top panels.

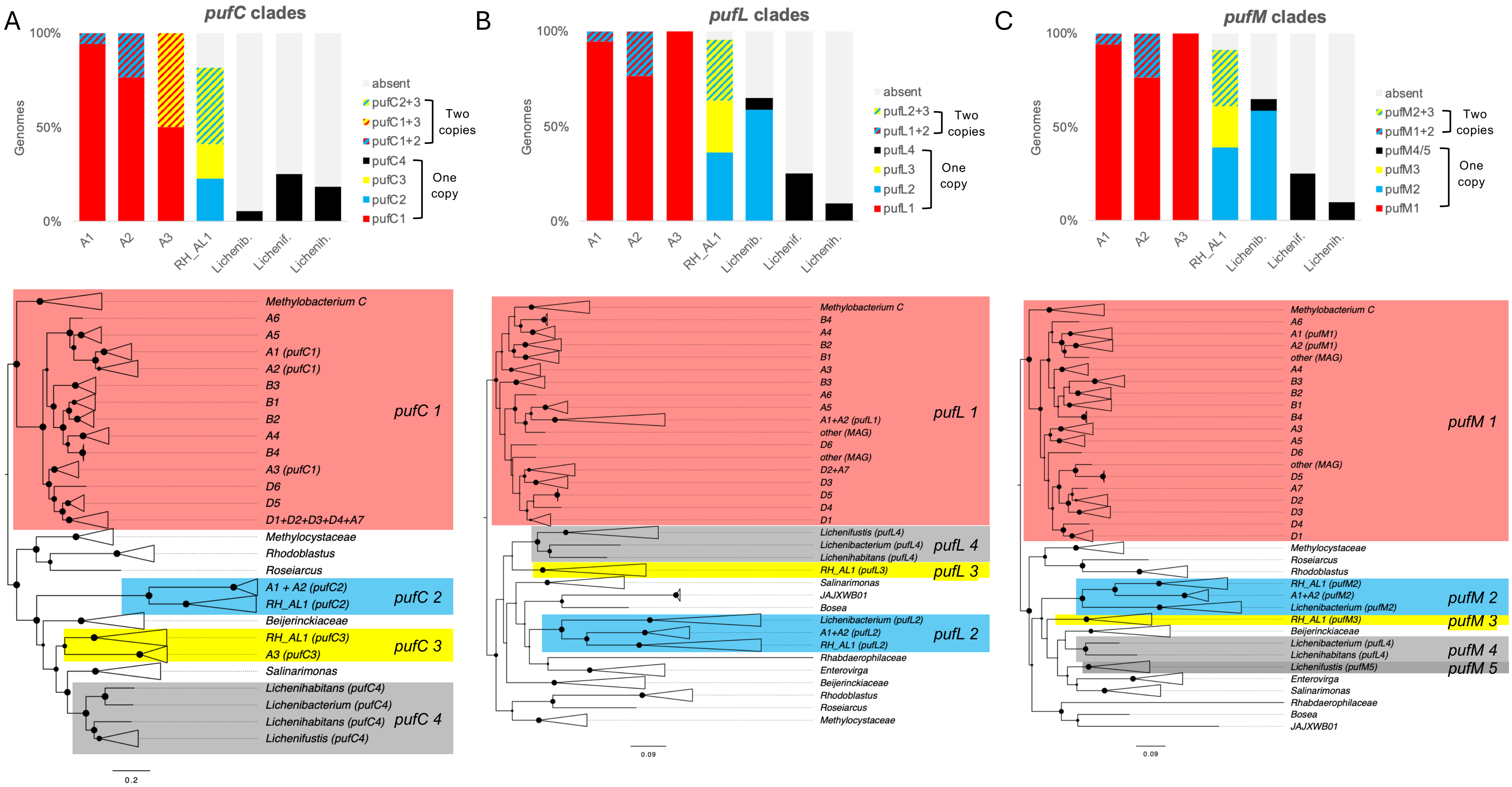

A
pufC clades
Genomes
100%
50%
0%
A1
A2
A3
RH_AL1
Lichenib.
Lichenif.
Lichenih.
absent
pufC2+3
pufC1+3
pufC1+2
Two copies
pufC4
pufC3
pufC2
pufC1
One copy
B
pufL clades
absent
pufL2+3
pufL1+2
Two copies
pufL4
pufL3
pufL2
pufL1
One copy
C
pufM clades
absent
pufM2+3
pufM1+2
Two copies
pufM4/5
pufM3
pufM2
pufM1
One copy
Methylobacterium C
A6
A5
A1 (pufC1)
A2 (pufC1)
B3
B1
B2
A4
B4
A3 (pufC1)
D6
D5
D1+D2+D3+D4+A7
pufC 1
Methylocystaceae
Rhodoblastus
Roseiarcus
A1 + A2 (pufC2)
RH_AL1 (pufC2)
pufC 2
Beijerinckiaceae
RH_AL1 (pufC3)
A3 (pufC3)
pufC 3
Salinarimonas
Lichenihabitans (pufC4)
Lichenibacterium (pufC4)
Lichenihabitans (pufC4)
Lichenifustis (pufC4)
pufC 4
0.2
Methylobacterium C
B4
A4
B2
B1
A3
B3
A6
A5
A1+A2 (pufL1)
other (MAG)
D6
other (MAG)
D2+A7
D3
D5
D4
D1
pufL 1
Lichenifustis (pufL4)
Lichenibacterium (pufL4)
Lichenihabitans (pufL4)
pufL 4
RH_AL1 (pufL3)
pufL 3
Salinarimonas
JAJXWB01
Bosea
Lichenibacterium (pufL2)
A1+A2 (pufL2)
RH_AL1 (pufL2)
pufL 2
Rhabdaerophilaceae
Enterovirga
Beijerinckiaceae
Rhodoblastus
Roseiarcus
Methylocystaceae
0.09
Methylobacterium C
A6
A1 (pufM1)
A2 (pufM1)
other (MAG)
A4
B3
B2
B1
B4
A3
A5
D6
other (MAG)
D5
A7
D2
D3
D4
D1
pufM 1
Methylocystaceae
Roseiarcus
Rhodoblastus
RH_AL1 (pufM2)
A1+A2 (pufM2)
Lichenibacterium (pufM2)
pufM 2
RH_AL1 (pufM3)
pufM 3
Beijerinckiaceae
Lichenibacterium (pufL4)
Lichenihabitans (pufL4)
pufM 4
Lichenifustis (pufM5)
pufM 5
Enterovirga
Salinarimonas
Rhabdaerophilaceae
Bosea
JAJXWB01
0.09
